# Structural mechanism of CARD8 regulation by DPP9

**DOI:** 10.1101/2021.01.13.426575

**Authors:** Humayun Sharif, L. Robert Hollingsworth, Andrew R. Griswold, Jeffrey C. Hsiao, Qinghui Wang, Daniel A. Bachovchin, Hao Wu

**Affiliations:** Department of Biological Chemistry and Molecular Pharmacology, Harvard Medical School, Boston, MA 02115, USA; Program in Cellular and Molecular Medicine, Boston Children’s Hospital, Boston, MA 02115, USA; Program in Biological and Biomedical Sciences, Harvard Medical School, Boston, MA 02115, USA; Weill Cornell/Rockefeller/Sloan Kettering Tri-Institutional MD-PhD Program, New York, NY, USA; Pharmacology Program, Weill Cornell Graduate School of Medical Sciences, Memorial Sloan Kettering Cancer Center, New York, New York 10065, USA; Chemical Biology Program, Memorial Sloan Kettering Cancer Center, New York, NY, USA

**Keywords:** CARD8, NLRP1, DPP9, inflammasome, pyroptosis, Val-boroPro (VbP), cryo-EM

## Abstract

CARD8 is a germline-encoded pattern recognition receptor that detects intracellular danger signals. Like the related inflammasome sensor NLRP1, CARD8 undergoes constitutive autoprocessing within its function-to-find domain (FIIND), generating two polypeptides that stay associated and autoinhibited. Certain pathogen- and danger-associated activities, including the inhibition of the serine dipeptidases DPP8 and DPP9 (DPP8/9), induce the proteasome-mediated degradation of the N-terminal (NT) fragment, releasing the C-terminal (CT) fragment to form a caspase-1 activating inflammasome. DPP8/9 also bind directly to the CARD8 FIIND, but the role that this interaction plays in CARD8 inflammasome regulation is not yet understood. Here, we solved several cryo-EM structures of CARD8 bound to DPP9, with or without the DPP inhibitor Val-boroPro (VbP), which revealed a ternary complex composed of one DPP9, the full-length CARD8, and one CARD8-CT. Through structure-guided biochemical and cellular experiments, we demonstrated that DPP9’s structure restrains CARD8-CT after proteasomal degradation. Moreover, although DPP inhibitors do not directly displace CARD8 from DPP9 *in vitro*, we show that they can nevertheless destabilize this complex in cells. Overall, these results demonstrate that DPP8/9 inhibitors cause CARD8 inflammasome activation via at least two distinct mechanisms, one upstream and one downstream of the proteasome.

## INTRODUCTION

Canonical inflammasomes are cytosolic supramolecular signaling complexes that form in response to diverse pathogenic and endogenous danger signals to induce pyroptotic cell death and cytokine activation (Broz and Dixit; Martinon et al., 2002; Shen et al., 2019). Human CARD8 was recently identified to assemble one such inflammasome (Johnson et al., 2018), but unlike most other inflammasome sensors, CARD8 is not a member of the nucleotide-binding domain and leucine-rich repeat containing (NLR) family. It instead has an unusual function-to-find domain (FIIND) at the core of the protein flanked by a disordered N-terminal region that varies between isoforms (Bagnall et al., 2008) and a C-terminal caspase activation and recruitment domain (CARD) (Figure 1A). The FIIND and CARD of CARD8 share domain organization with those of one NLR inflammasome sensor, NLRP1. For both proteins, the FIIND possesses autoproteolytic activity which generates a noncovalent complex of the repressive N-terminal fragment (NT) that includes the ZU5 subdomain of the FIIND, and the inflammatory C-terminal fragment (CT) that includes the UPA subdomain of the FIIND and the CARD (UPA-CARD) (D’Osualdo et al., 2011; Finger et al., 2012) (Figure 1A). The inflammatory CT must be released from the repressive NT for CARD8 activation (Johnson et al., 2018).

**Figure 1.**
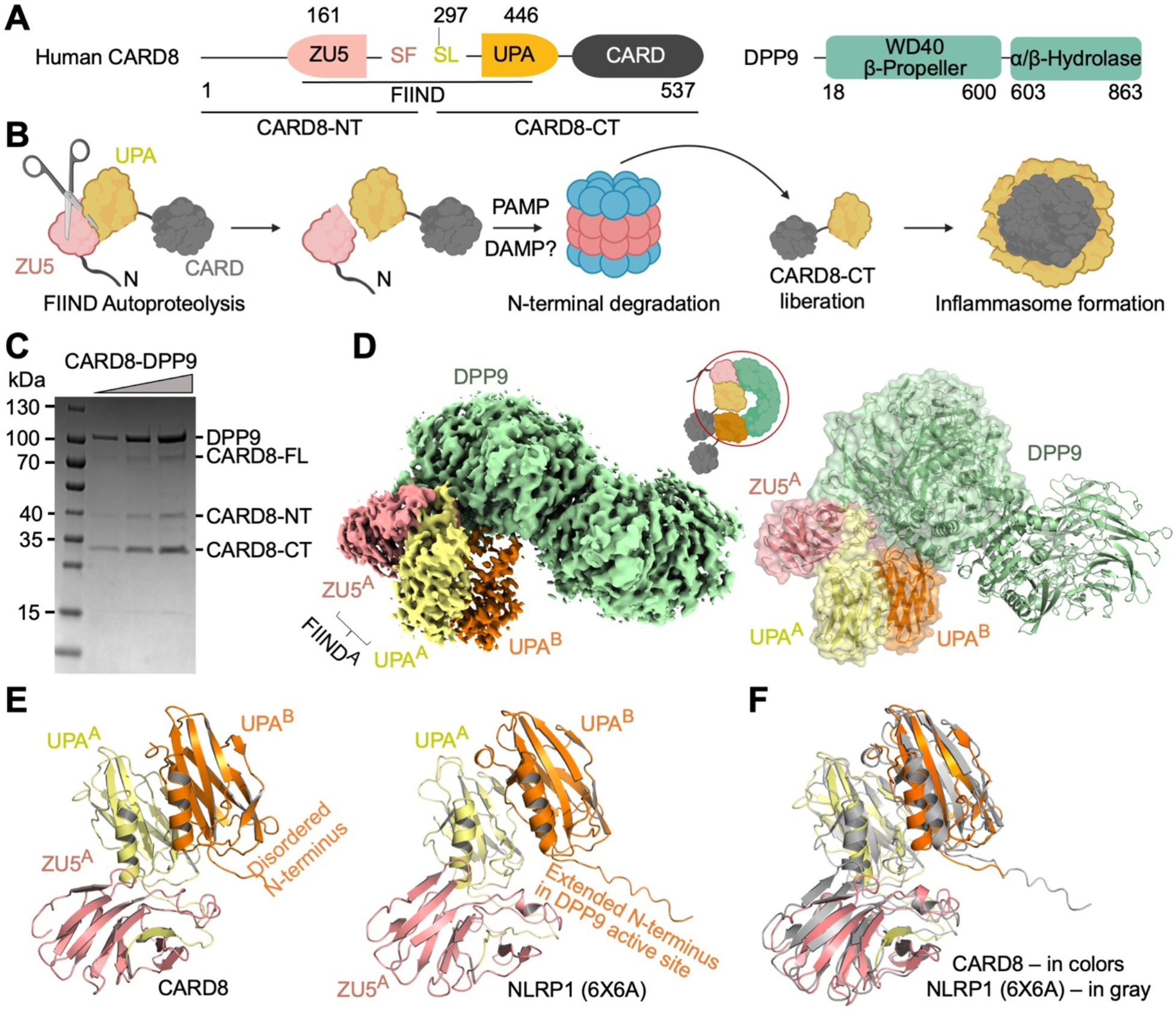
Cryo-EM Structure of the CARD8-DPP9 Complex. (A) Domain architecture of CARD8 and DPP9. The FIIND autoprocessing site is between F296 and S297. (B) Schematic diagram of CARD8 activation by N-terminal degradation. (C) SDS-PAGE of the purified CARD8-DPP9 complex, revealing both processed and unprocessed CARD8. The CARD8-CT appears overstochiometric to the CARD8-NT. (D) Cryo-EM density (left) and model (right) of the CARD8-DPP9 complex at a 2:2 CARD8-DPP9 stoichiometry colored as in (A). One complete ZU5-UPA pair (site A) and a second UPA (site B) associate with a DPP9 monomer. A schematic (middle) denotes the construct versus the ordered, resolved portions of the protein (red circle). (E) Comparison of the CARD8 (left) and NLRP1 (right) ZU5^A^-UPA^A^-UPA^B^ modules. The N-terminus for NLRP1-UPA^B^ extends to bind into the DPP9 active site whereas the same segment is disordered for CARD8-UPA^B^. (F) Orientational difference between CARD8 (colored) and NLRP1 (grey). DPP9 dimers in the CARD8-DPP9 and NLRP1-DPP9 (6X6A) complexes were superimposed, showing the different relative relationships between the domains. See also Figure S1 and S2, and Table S1.

DPP8/9 inhibitors, including VbP, activate the CARD8 inflammasome in human macrophages and resting lymphocytes and the NLRP1 inflammasome in skin and airway epithelial cells (Johnson et al., 2020; Johnson et al., 2018; Linder et al., 2020; Robinson et al., 2020; Zhong et al., 2018). The mechanisms of DPP8/9 inhibitor-induced CARD8 and NLRP1 inflammasome activation have not yet been fully established. We recently discovered that DPP8/9 inhibitors induce the proteasomal degradation of the NT fragments (Chui et al., 2020; Chui et al., 2019) releasing the noncovalently associated CT fragments from autoinhibition (Figure 1B). However, the identity and activation mechanism of the key E3 ligase remains unknown. In addition, CARD8 and NLRP1 directly bind to DPP8 and DPP9 (Griswold et al., 2019; Zhong et al., 2018). Intriguingly, structural and biochemical studies recently revealed that DPP9 binding restrains NLRP1 activation by sequestering the freed NLRP1-CT fragment, and VbP activates human NLRP1, at least in part, by disrupting this interaction (Hollingsworth et al., 2020b; Huang et al., 2020). In contrast, VbP does not directly disrupt the CARD8-DPP9 interaction (Griswold et al., 2019), indicating significant differences between the NLRP1 and CARD8 regulatory mechanisms.

Here, we investigated the structural and cellular mechanisms of CARD8 regulation by DPP9 and DPP8/9 inhibitors, revealing interesting differences from the recent complementary findings for NLRP1 (Hollingsworth et al., 2020b; Huang et al., 2020). Most notably, we discovered that DPP9 binding also restrains the free CARD8-CT, but the CARD8-CT, unlike the NLRP1-CT, does not closely interact with the DPP9 active site, explaining why DPP8/9 inhibitors do not directly displace the CARD8-CT. However, we discovered that DPP8/9 inhibitors do destabilize this interaction in cells, showing that these compounds nevertheless induce CARD8 inflammasome activation via two separate mechanisms.

## RESULTS

### Cryo-EM Structure of the CARD8^A^-CARD8^B^-DPP9 Ternary Complex

We determined the cryo-EM structure of the co-expressed CARD8-DPP9 complex (Figure 1C) at a resolution of 3.3 Å (Figure S1, Table S1). The overall cryo-EM structure revealed 2 copies (molecules A and B) of CARD8 bound to a single DPP9 subunit, forming a CARD8^A^-CARD8^B^-DPP9 ternary complex (Figure 1D). Like NLRP1 (Hollingsworth et al., 2020b; Huang et al., 2020), molecule A has a full-length (FL) FIIND composed of the ZU5-UPA pair, and molecule B contains only the UPA. Other domains of CARD8 were not visible in the EM density, likely due to their flexible linkage to the FIIND and their intrinsically disordered sequence composition (Chui et al., 2020). For FIIND^A^, there is weak electron density around the autoprocessing site; given that SDS-PAGE suggested over half of CARD8 was autoprocessed (Figure 1C), we built the atomic model as an autoprocessed FIIND although FIIND^A^ is likely a mixture of unprocessed and processed FIIND. Unlike the NLRP1-DPP9 complex structure in which both monomers of the DPP9 dimer are each bound to 2 copies of NLRP1 (4:2 NLRP1:DPP9 stoichiometry) (Hollingsworth et al., 2020b), the final CARD8-DPP9 map has a 2:2 CARD8:DPP9 stoichiometry in which one DPP9 monomer of the DPP9 dimer is not bound to CARD8. This result is explained by the overabundance of the 2:2 DPP9:CARD8 complex particles (Figure S1C) over 4:2 DPP9:CARD8 complex particles (Figure S1D), likely reflecting understoichiometric CARD8 expression relative to DPP9 but with no apparent functional significance.

A striking difference between the CARD8-DPP9 and the NLRP1-DPP9 structures exists at the N-terminus of UPA^B^, despite the 50% sequence identity between CARD8 and NLRP1 FIINDs and the similar structural architecture of the complexes (Figure 1E, S2A-B). Unlike in NLRP1 (Hollingsworth et al., 2020b), the N-terminus of CARD8-UPA^B^ does not insert into the DPP9 active site and is largely disordered (Figure 1E). In addition, when DPP9 monomers are aligned, the bound FIIND^A^-UPA^B^ complex in CARD8 exhibits a significant shift relative to that in NLRP1, ~9° in rotation and ~1Å in translation (Figure 1F), suggesting some differences in the exact binding mode to accommodate the sequence divergence. Among other ZU5 and UPA fold-containing proteins, the FIIND β-sandwich folds are similar to those of ankyrin-B (Wang et al., 2012) and the axonal guidance receptor Unc5b (Wang et al., 2009) with two notable exceptions. First, neither ankyrin-B nor Unc5b contains an UPA-UPA interaction (Wang et al., 2012; Wang et al., 2009) (Figure S2C). Second, the FIIND^A^ module largely recapitulates the spatial relationship of ZU5 and UPA in Unc5b, but for ankyrin-B, which contains a covalently linked ZU5(1)-ZU5(2)-UPA module, the FIIND^A^ module instead mimics the arrangement of ankyrin ZU5(1) and UPA rather than the covalently linked ZU5(2)-UPA relationship (Figure S2C).

### CARD8^B^ Is N-terminally Degraded and Its Binding Is Not Affected by DPP9 Inhibitors in Vitro

We noticed a larger amount of CARD8-CT than CARD8-NT in co-eluted fraction of the CARD8-DPP9 complex (Figure 1C), suggesting that the UPA^B^ did not have an associated ZU5^B^. This observation could be explained by N-terminal degradation of CARD8 (Chui et al., 2019; Johnson et al., 2018; Sandstrom et al., 2019) during normal protein turnover, which would free the noncovalently associated CT for ternary complex formation with FIIND^A^ and DPP9. Indeed, if we modeled the theoretical ZU5^B^ subdomain based on rigid superimposition of FIIND^A^ (containing both ZU5^A^ and UPA^A^) onto UPA^B^, steric clashes would have existed between this ZU5^B^ and both DPP9 and FIIND^A^ (Figure 2A). To assess whether autoproteolysis and presumably concomitant N-terminal degradation of the CARD8-NT was required for site B binding, we solved the structure of autoproteolytic-deficient CARD8-S297A in complex with DPP9 (Figure S3, Table S1). The density of UPA^B^ was entirely missing in this structure (Figure 2B). Together, these data suggest that each DPP9 monomer binds 1 molecule of FL-CARD8 and 1 molecule of already N-terminally degraded CARD8-CT.

**Figure 2.**
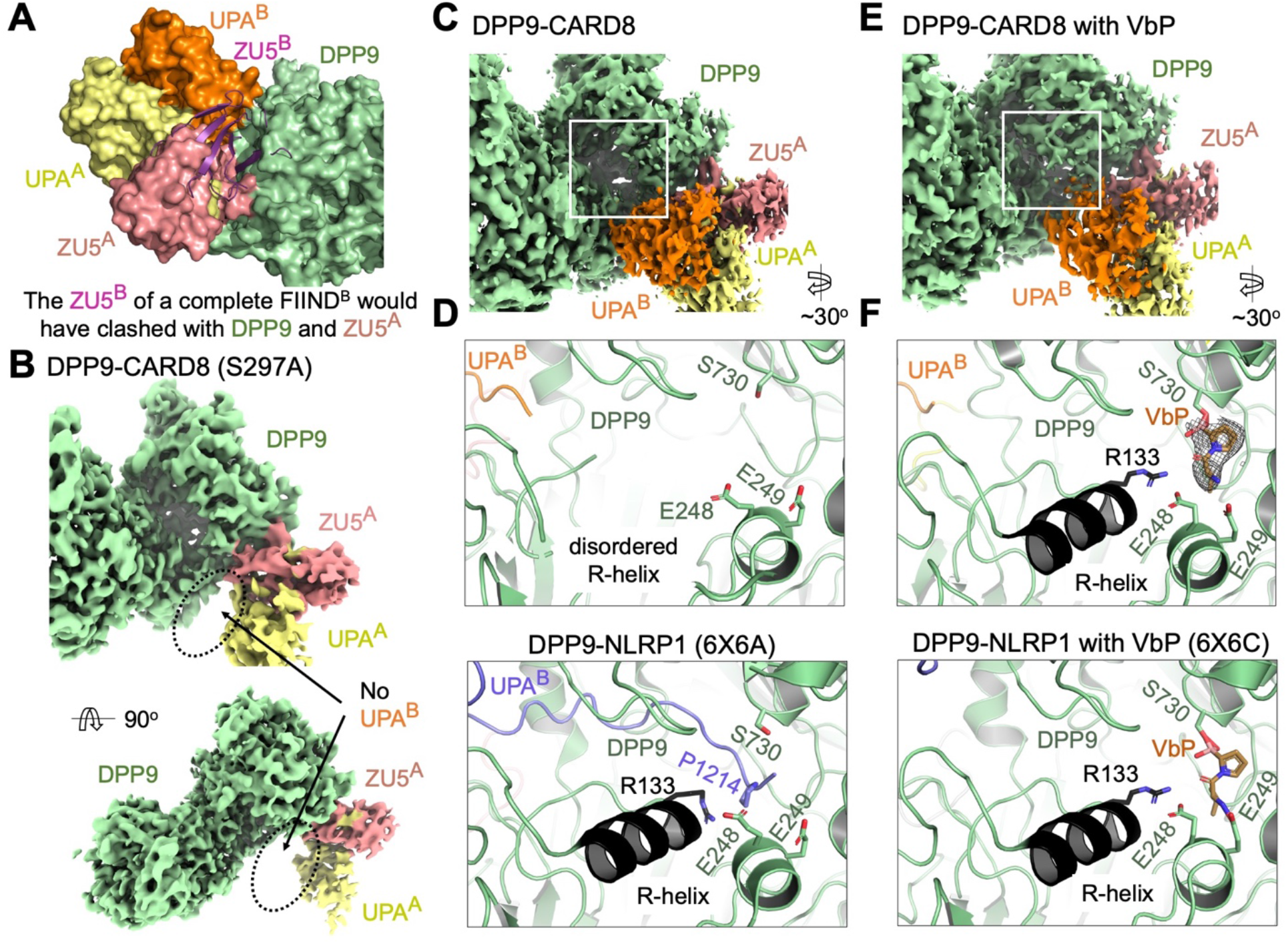
Binding of CARD8-CT at Site B Which is Not Disrupted by DPP9 Inhibitors. (A) Superimposition of the ZU5^A^-UPA^A^ relationship onto UPA^B^ to generate a theoretical ZU5^B^ (purple). A theoretical ZU5^B^ cannot be accommodated by the structure, explaining why site B is a dissociated UPA^B^, rather than a complete FIIND. (B) Cryo-EM density of the DPP9-CARD8 (S297A) complex with no visible UPA^B^ density. Autoproteolysis, and presumably, N-terminal degradation is required for site B association. (C) Density of the DPP9-CARD8 complex near the DPP9 active site. (D) Zoom-in of the DPP9 substrate tunnel in the DPP9-CARD8 complex. CARD8-UPA^B^ binds near, but not into, the DPP9 substrate tunnel and the R-helix remains disordered. In comparison, zoom-in view of the NLRP1-DPP9 complex (PDB ID: 6X6A) at the DPP9 active site shows occupancy by the NLRP1-CT peptide. (E) Density of the DPP9-CARD8 complex with bound VbP near the active site. (F) Zoom-in of the DPP9 substrate tunnel in the DPP9-CARD8 complex with VbP. VbP binds near the active site, causing R-helix ordering, but does not displace CARD8-UPA^B^. in comparison, zoom-in view of the NLRP1-DPP9-VbP complex (PDB ID: 6X6C) at the DPP9 active site shows that VbP displaces the NLRP1-CT peptide. See also Figure S3 and S4, and Table S1.

The CARD8-CT binds adjacent to, but not inside of, the DPP9 substrate tunnel leading to the active site (Figure 2C-D). The DPP9 active site is known to undergo substantial rearrangement at a loop segment that folds into an α-helix, known as the R-helix, when R133 of the R-helix engages a substrate (Ross et al., 2018). Indeed, the R-helix is disordered in the CARD8-DPP9 complex (Figure 2C-D). This is directly in contrast to NLRP1, which buries the N-terminus of NLRP1-UPA^B^ into the DPP9 active site, resulting in R-helix ordering (Figure 2D) (Hollingsworth et al., 2020b). The competitive DPP9 inhibitor VbP also causes R-helix ordering (Ross et al., 2018) and activates CARD8 (Johnson et al., 2018). To investigate whether inhibitor binding affected the UPA^B^ CARD8-CT, we solved the CARD8-DPP9-VbP structure at 3.3 Å resolution (Figure S4, Table S1). VbP bound directly into the DPP9 active site and resulted in R-helix ordering but did not displace either CARD8 molecule (Figure 2E-F, Figure S4G). This finding was again in direct contrast to human NLRP1, as VbP almost completely displaced NLRP1-UPA^B^ from DPP9 (Figure 2F) (Hollingsworth et al., 2020b). Collectively, these findings rationalize the discrepancies observed in co-immunoprecipitation experiments between the human NLRP1-DPP9 complex and the CARD8-DPP9 complex (Griswold et al., 2019; Zhong et al., 2018): human NLRP1-CT binds into the DPP9 active site and this interface is disrupted by inhibitors such as VbP, whereas CARD8-CT does not bind into the DPP9 active site and its binding is not directly affected by VbP.

### Three Interfaces Mediate DPP9-Ternary Complex Assembly

Like NLRP1, three large interfaces are involved in CARD8-DPP9 association: (I) ZU5^A^-DPP9 (~950 Å^2^/partner), (II) UPA^B^-DPP9 (~600 Å^2^/partner), and (III) UPA^A^-UPA^B^ (800 Å^2^/partner) (Figure 3A). Interface I is formed by β9 and the β4-β5 loop in ZU5^A^ and the WD40 domain of DPP9. Hydrophobic residues V277 and L278 in ZU5^A^ pack against LLL100 in DPP9 and K272-E274 in ZU5^A^ buries significant surface area by DPP9 association. Several notable hydrogen bonding interactions include M276 and E279 in ZU5^A^ between L102 and R96 in DPP9, respectively. Interface II involves interactions near the DPP9 active site tunnel including Y322 near the UPA^B^ N-terminus with DPP9’s K43. Additionally, H39 in DPP9 wedges between T430 and E431 in UPA^B^ with hydrogen bonding interactions between both. Because the UPA^B^ N-terminus is disordered, interface II between CARD8 and DPP9 is far less extensive compared to interface II in the NLRP1-DPP9 complex. Interface III, on the other hand, is very similar to NLRP1 in that the sides of the UPA^A^ and UPA^B^ β-sandwich folds closely associate. Notably, F405/Y406 on UPA^A^ (IIIa) and L368/F370 on UPA^B^ (IIIb) bury significant surface area through association (Figure 3A).

**Figure 3.**
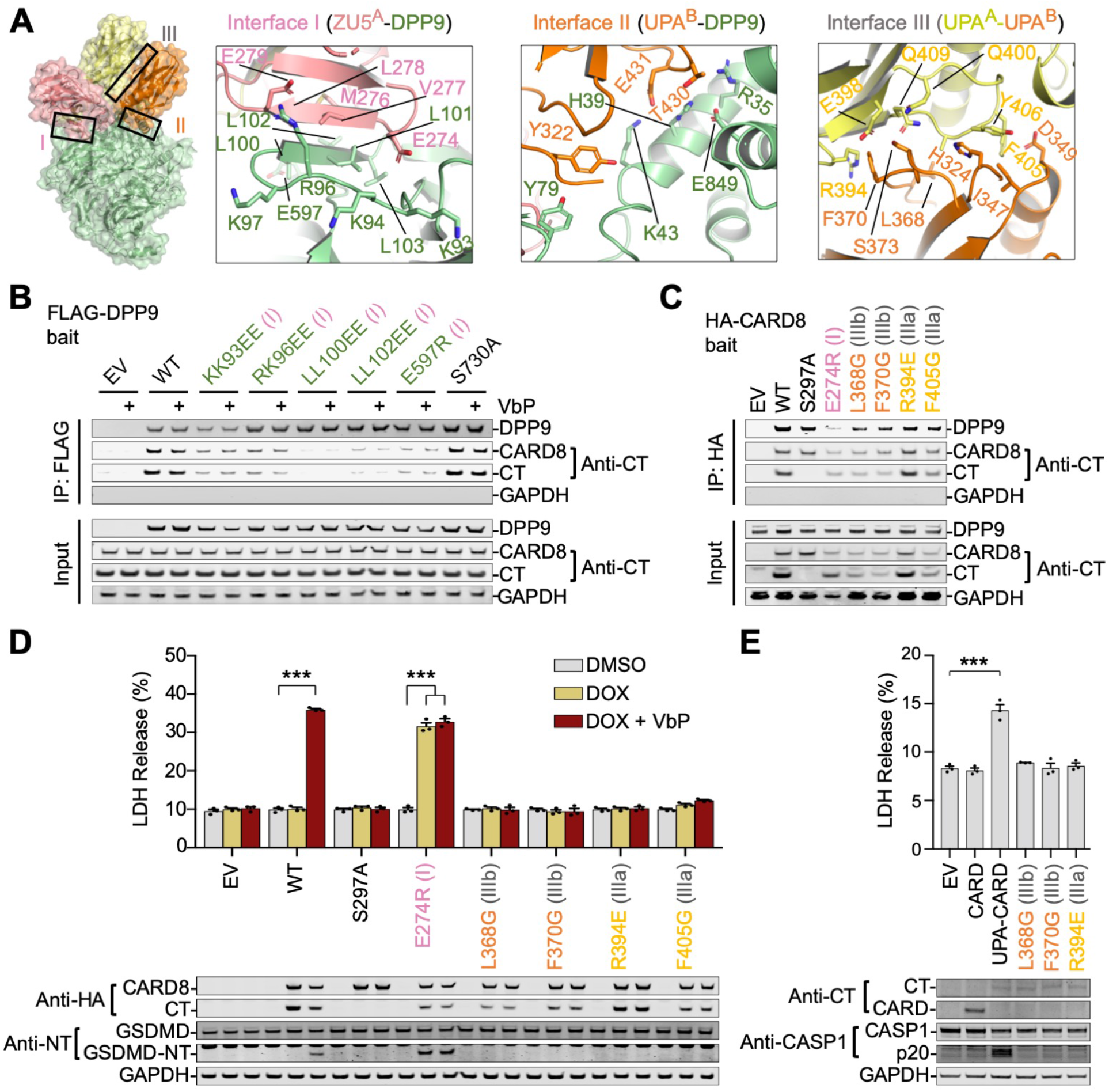
Mutational Analyses on the CARD8-DPP9 Interaction. (A) Overview of interfaces I (Zu5^A^-DPP9), II (UPA^B^-DPP9), and III (UPA^A^-UPA^B^) that mediate complex assembly. (B) FLAG co-immunoprecipitation of His-tagged CARD8 by structure-guided FLAG-tagged DPP9 mutants in the presence and absence of VbP. Constructs were expressed separately in *DPP8/9 DKO* HEK 293T cells and VbP was added to the combined lysates. Immunoblots are representative of 2 independent experiments. (C) HA co-immunoprecipitation of DPP9 by structure-guided HA-tagged CARD8 mutants from THP-1 CARD8^-/-^ cells. The interface I mutant disrupted CARD8-DPP9 association, but the interface III mutants and the CARD8 autoprocessing-deficient mutant (S297A) did not. Immunoblots are representative of 2 independent experiments. (D) Cell death in THP-1 CARD8^-/-^ monocytes stably reconstituted with the indicated Tet-inducible WT and mutant FL-CARD8 constructs. Interface I mutations led to CARD8 autoactivation while interface III mutations abolished CARD8 activity as measured by LDH release (top) and Western blot (bottom) of inflammasome components. Mean ± SEM are shown from 3 independent biological replicates. *** p < 0.001 by 2-way ANOVA with Tukey’s multiple comparison correction. (E) Cell death in HEK 293T cells that stably express GSDMD and CASP-1 by transient expression of indicated CARD8-CT constructs. UPA-CARD enhanced while Interface III mutations abolished inflammasome activity as measured by LDH release (top) and Western blot (bottom) of inflammasome components. Mean ± SEM are shown from 3 independent biological replicates. *** p < 0.001 by Student’s two-sided T-test.

Structure-guided DPP9 mutations on interface I reduced the CARD8 interaction (Figure 3B) without affecting DPP9’s activity (Hollingsworth et al., 2020b). Addition of VbP during coimmunoprecipitation did not reduce the DPP9-CARD8 interaction, underscoring the different binding modes of CARD8-UPA^B^ and VbP (Figure 3B). Autoproteolysis deficient CARD8-S297A was able to bind DPP9 at near wild-type (WT) levels (Figure 3C), indicating that the DPP9-CARD8^A^ interaction does not require CARD8^B^.

### DPP9 Binding Restrains the CARD8 Inflammasome and UPA-UPA Interactions are Necessary for Inflammasome Signaling

We next generated structure-guided mutations on CARD8 to assess the role of the DPP9-ternary complex interfaces on inflammasome regulation. While most mutations abolished FIIND autoproteolysis, we managed to identify mutations on interface I (E274R of ZU5^A^), IIIa (R394E and F405G of UPA^A^), and IIIb (L368G and F370G of UPA^B^) that preserved autoproteolysis as evidenced by similar levels of CARD8-CT in mutant and WT samples (Figure 3C). As expected, the interface I mutants abolished DPP9 binding. In contrast, interface III mutants maintained DPP9 binding (Figure 3C). This further illustrates differences between NLRP1 and CARD8 in their assembly of DPP9-ternary complexes: for CARD8, FIIND^A^ can bind without an associated UPA^B^. NLRP1, on the other hand, requires both copies for cooperative DPP9 association (Hollingsworth et al., 2020b).

We next assessed the functional impacts of these mutations by stably reconstituting CARD8^-/-^ THP-1 cells (Johnson et al., 2018) with Tet-inducible CARD8 constructs. For WT CARD8, the combination of doxycycline and VbP were required for cell death and GSDMD cleavage (Figure 3D). Doxycycline-induced expression of CARD8-E274R, an interface I mutant that cannot bind DPP9, caused pyroptosis in the absence of VbP with no significant increase upon its addition (Figure 3D). Thus, DPP9 binding restrains CARD8 inflammasome activation.

Despite maintaining FIIND autoprocessing, interface III mutations completely abolished inflammasome activity, even in the presence of the activating ligand VbP (Figure 3D). We reasoned that the observed UPA-UPA interface might be preserved on the inflammasome filament, as the UPA itself promotes CARD oligomerization and inflammasome activity (Hollingsworth et al., 2020a; Qin et al., 2020). Additionally, NLRP1 interface III mutations abolished inflammasome signaling in cells (Hollingsworth et al., 2020b) and disrupted filament formation in vitro (Huang et al., 2020). Consistent with this premise, only direct expression of the WT UPA-CARD, but not UPA-CARDs with interface III mutations nor the CARD alone, resulted in robust inflammasome signaling in a reconstituted HEK 293T system (Figure 3E).

### DPP9 Requires FL-CARD8 to Sequester and Repress CARD8-CT

The above results indicate that CARD8-DPP9 binding is required to prevent the formation of an UPA-CARD inflammasome. We next asked whether FIIND^A^ was required to load CARD8-CT onto DPP9. We co-expressed CARD8-FIIND-S297A (FIIND-SA), which can only bind DPP9 site A, with CARD8-CT-FLAG, which can only bind DPP9 site B in HEK 293T cells. FLAG immunoprecipitation indicated that CARD8-CT only captures DPP9 in the presence of FIIND-SA (Figure 4A), revealing that the CARD8-CT requires non-degraded FIIND^A^ for DPP9 association. These data are consistent with the reduction of both CARD8 and CARD8-CT interaction by structure-guided DPP9 mutations on interface I (Figure 3B). FIIND-SA with a site I mutation that abolishes DPP9 binding still associates with CARD8-CT (Figure 4A), indicating that CARD8 dimerization can occur in the absence of DPP9 through interface III. Mutations on interface III of FIIND-SA reduced its interaction with CARD8-CT as well as DPP9 (Figure 4A).

**Figure 4.**
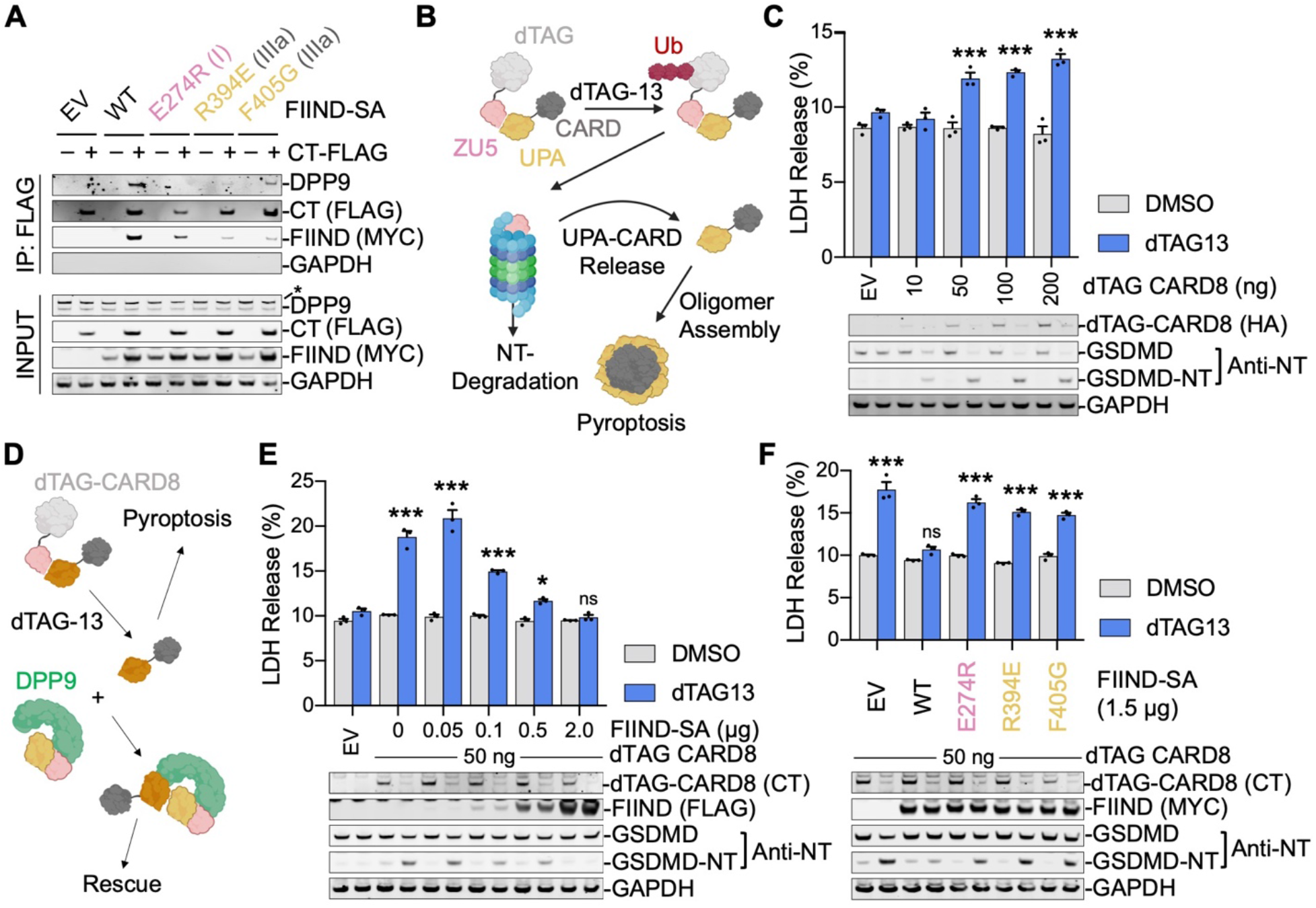
Suppression of CARD8-CT at Site B of DPP9 by Binding of FL-CARD8 to Site A. (A) FLAG co-immunoprecipitation of FLAG-tagged CARD8-CT with the indicated EV (empty vector), WT and mutant CARD8-FIIND-S297A (FIIND-SA) constructs co-expressed in HEK 293T cells. The interface I mutation preserved CARD8^A^-CARD8^B^ interaction while abolishing DPP9 binding, and interface III mutations on FIIND-SA impaired CARD8^A^-CARD8^B^ interaction and the DPP9 interaction. Immunoblots are representative of 2 independent experiments. (B) Schematic of the dTAG experimental system. FL-CARD8 is fused to N-terminal FKBP12^F36V^ (dTAG-CARD8), which mediates ubiquitination and degradation of the fusion protein in the presence of the small molecule degrader dTAG-13. (C) Cell death induced by dTAG13 (500 nM, 3h) via N-terminal degradation in a reconstituted HEK 293T system expressing an increasing amount of dTAG-CARD8 (10-200 ng transfected plasmid). Pyroptosis was measured by LDH release (top) and GSDMD cleavage (bottom). Mean ± SEM are shown from 3 independent biological replicates. *** p < 0.001 denotes comparison to EV treated with DMSO by 2-way ANOVA with Tukey’s multiple comparison correction. (D) Schematic of rescuing dTAG-13 dependent pyroptosis with the DPP9-ternary complex. (E) Rescue of dTAG-13-induced dTAG-CARD8-mediated cell death in a HEK 293T system by increased co-expression of FIIND-SA. Pyroptosis was measured by LDH release (top) and GSDMD cleavage (bottom). dTAG-13 was used at 500 nM for 3 hr. Mean ± SEM are shown from 3 independent biological replicates. ***, * and ns denote p < 0.001, p < 0.1 and not significant, respectively, in comparison to EV treated with DMSO by 2-way ANOVA with Tukey’s multiple comparison correction. (F) Failed rescue of CARD8-CT-mediated cell death by mutant FIIND-SA constructs that abolish tertiary complex formation, measured by LDH release (top) and GSDMD processing (bottom). Reconstituted HEK 293T cells expressing dTAG-CARD8 and FIIND-SA were treated with dTAG-13 (500 nM, 3h). Mean ± SEM are shown from 3 independent biological replicates. *** and ns denote p < 0.001 and not significant, respectively, in comparison to EV treated with DMSO by 2-way ANOVA with Tukey’s multiple comparison correction.

Since DPP9 association is required to repress CARD8 inflammasome activity and because CARD8-CT requires FIIND^A^ for DPP9 association, we postulated that DPP9-ternary complex formation is required to repress CARD8-CT inflammasome activity. To test this hypothesis, we leveraged the dTAG degron tag system, which facilitates controlled degradation of any FKBP12^F36V^ fusion protein in response to the ligand dTAG-13 (Nabet et al., 2018), to modulate the amount of free CARD8-CT. Briefly, when fused N-terminally to CARD8 (dTAG-CARD8-FL), dTAG-13 induces degradation of the CARD8-NT and liberates the free CARD8-CT due to FIIND autoprocessing (Figure 4B). Indeed, transient expression of dTAG-CARD8-FL in a reconstituted HEK 293T cell system caused marked cell death upon stimulation with dTAG-13 (Figure 4C). In our system, titration of the dTAG-CARD8-FL plasmid modulated the level of cell death in response to dTAG-13, as demonstrated by LDH release.

We next co-expressed increasing amounts of FIIND-SA along with dTAG-CARD8-FL in the reconstituted HEK 293T cell system (Figure 4D). Indeed, FIIND-SA rescued dTAG-13-induced cell death in a dose-dependent manner (Figure 4E). Additionally, interface I and III mutations abrogated FIIND-SA rescue (Figure 4F). Notably, the interface I mutant still preserved FIIND^A^-UPA^B^ association through interface III (Figure 4A), demonstrating that DPP9 binding, and not CARD8 dimerization, represses the activity of CARD8-CT. Thus, FL-CARD8 recruits CARD8-CT to the DPP9-ternary complex, which ultimately represses CARD8 inflammasome formation.

### VbP Activates CARD8 by N-terminal Degradation and DPP9-Ternary Complex Disruption

The above results indicate that DPP9 binding is imperative to its regulation of the CARD8 inflammasome. However, DPP9’s catalytic activity also contributes to inflammasome regulation. DPP9 cleaves two residues from the N-terminus of peptides that contain a penultimate proline or alanine (NH2-X-P or NH2-X-A) (Geiss-Friedlander et al., 2009; Olsen and Wagtmann, 2002). The N-terminus of UPA of NLRP1, but not CARD8, has such a motif (Figure 1A). We recently demonstrated that inhibition of cellular DPP catalytic activity with VbP stimulates N-terminal degradation of CARD8 through a Cullin E3 ubiquitin ligase pathway (Chui et al., 2020). Thus, there are likely at least two distinct checkpoints regulating CARD8 activity: one upstream of the proteasome through CARD8 N-terminal degradation, and the other downstream of the proteasome through regulation of the DPP9-ternary complex.

We therefore sought to explain how DPP9 represses the activity of CARD8 without cleaving it directly. While VbP treatment causes some CARD8 degradation, it can take many hours to detect (Chui et al., 2019; Griswold et al., 2019). These kinetics are inconsistent with the rapid pyroptosis observed in macrophages and lymphocytes following DPP8/9 inhibition (Johnson et al., 2020; Johnson et al., 2018), which prompted us to ask whether VbP also affects the DPP9-CARD8 ternary complex. While CARD8 still binds DPP9 in the presence of VbP (Figure 2B), we designed alternative assays to detect DPP9-ternary complex disruption *in vitro* and in cells. *In vitro*, we isolated the DPP9-ternary complexes on FLAG beads, treated the mixture with compounds, and looked for direct displacement as evident by release of either NLRP1-CT or CARD8-CT in the wash and the reduction of the bound amount in the elute (Figure 5A). As expected, NLRP1-CT is directly displaced by the structurally unrelated DPP9/8 inhibitors VbP and 8J (Van Goethem et al., 2008), shown by its release in the wash (Figure 5B). The amount of NLRP1-CT was also reduced in the elute from VbP and 8J-treated samples in comparison with those treated with the amino peptidase inhibitor bestatin methyl ester (MeBS), or the DMSO vehicle control. In contrast, DPP8/9 inhibitors did not increase the amount of CARD8-CT released from the beads in the wash or reduce the bound CARD8-CT in the elute, consistent with the premise that VbP itself does not directly displace CARD8 from DPP9 (Figure 5C).

**Figure 5.**
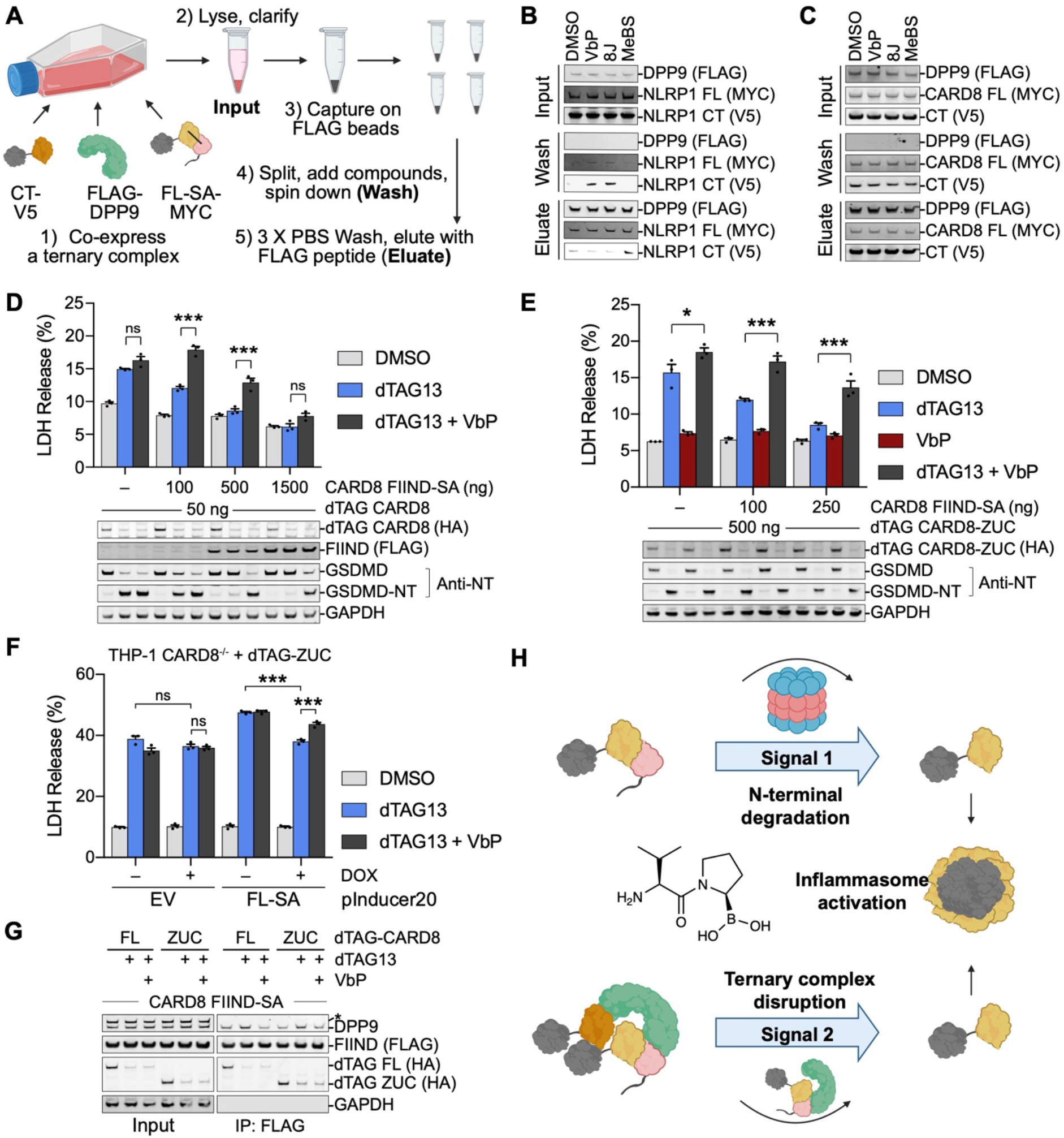
Two Distinct Ways to Activate the CARD8 Inflammasome. (A) Schematic of a direct displacement assay for DPP9 inhibition. Co-expressed FLAG-DPP9, NLRP1 or CARD8 CT-V5 and full-length NLRP1 or CARD8 SA mutant (FL-SA-MYC) (Input) were captured on FLAG beads. The bound components were displaced with DPP8/9 inhibitors or control compounds (Wash), and then eluted with 3x-FLAG peptide (Eluate). (B) Displacement of NLRP1-CT from DPP9 by the DPP9/8 inhibitors VbP and 8J in vitro as depicted in (A). The displacement was apparent in the Wash, and also visible in the Eluate. The vehicle control DMSO or the non-selective aminopeptidase inhibitor MeBs did not cause displacement. (C) Lack of displacement of CARD8-CT from DPP9 in vitro by any of the compounds. Immunoblots are representative of 2 independent experiments. (D) Further enhancement of cell death by VbP in dTAG13-induced CARD8 activation in the presence of FIIND-SA rescue. Reconstituted HEK 293T cells expressing dTAG-CARD8 and increasing amount of FIIND-SA were treated with dTAG-13 (500 nM) alone, or both dTAG-13 and VbP (10 μM) for 3 hr. Cell death was measured by LDH release (top) and GSDMD processing (bottom). The enhancement was most apparent at a lower dose of transfected FIIND-SA. Mean ± SEM are shown from 3 independent biological replicates. *** and ns denote p < 0.001 and not significant, respectively, by 2-way ANOVA with Tukey’s multiple comparison correction. (E) VbP disruption of FIIND-SA mediated cell death rescue independent of VbP-induced CARD8 degradation. Reconstituted HEK 293T cells were expressed with dTAG-CARD8-ZUC, which does not contain the N-terminal disordered sequence to undergo VbP-induced degradation, and increasing amount of FIIND-SA, and treated with dTAG-13 (500 nM), VbP (10 μM), or dTAG13 plus VbP for 3 hr. Mean ± SEM are shown from 3 independent biological replicates. * and *** denote p < 0.05 and p < 0.001, respectively, by 2-way ANOVA with Tukey’s multiple comparison correction. (F) Functional disruption of FL-SA rescue of dTAG13-induced CARD8 activation by VbP in monocytes. THP-1 CARD8^-/-^ cells were stably reconstituted with dTAG-ZUC, with or without the Tet-inducible FL-SA (S297A) construct. While doxycycline (DOX)-dependent FL-SA expression dampened dTAG-13 induced pyroptosis, VbP disrupted this rescue. Mean ± SEM are shown from 3 independent biological replicates. *** and ns denote p < 0.001 and not significant, respectively, by 2-way ANOVA with Tukey’s multiple comparison correction. (G) Reduction of co-immunoprecipitated DPP9 by FLAG-tagged CARD8-FIIND-S297A (FIIND-SA). HEK 293T cells were co-expressed with FLAG-FIIND-SA and dTAG-CARD8 (FL) or dTAG-CARD8-ZUC (ZUC) constructs and treated with dTAG13 (500 nM) alone or dTAG13 plus VbP (10 μM) for 24 hr. * indicates non-specific band. (H) Model of CARD8 inflammasome activation upstream of the proteosome (signal 1) and downstream of the proteasome (signal 2). VbP contributes to activation via both mechanisms.

It remained possible that VbP could disrupt the CARD8 DPP9-ternary complex in cells. In order to test this hypothesis, we designed a HEK 293T system that decoupled N-terminal degradation from DPP9-ternary complex formation. We generated DPP9-ternary complexes with dTAG-CARD8-FL, which was completely degraded to release CARD8-CT in the presence of dTAG-13, and FIIND-SA, which lacks a disordered N-terminal segment required for VbP-stimulated N-terminal degradation (Chui et al., 2020) (Figure 5D). As expected, FIIND-SA repressed CARD8 activity. Remarkably, however, co-treatment with both dTAG-13 and VbP increased cell death and GSDMD processing compared to dTAG-13 alone. To further increase confidence that this VbP-mediated effect was independent of additional dTAG-CARD8-FL degradation, we generated a CARD8 dTAG-ZU5-UPA-CARD (dTAG-CARD8-ZUC) construct which is resistant to VbP induced N-terminal degradation and does not respond to VbP alone in HEK 293T cells (Figure 5E). Once again, VbP increased dTAG-13-dependent pyroptosis most noticeably in the presence of FIIND-SA.

Next, we stably expressed this CARD8 dTAG-ZUC construct in CARD8^-/-^ THP-1 cells (Figure 5F). As expected, dTAG-13 induced rapid pyroptosis. Likewise, doxycycline-inducible CARD8-S297A-FL expression reduced sensitivity to dTAG-13, demonstrating that the DPP9-ternary complex can indeed repress CARD8 signaling in monocytes. Notably, VbP only synergized with dTAG-13 in the presence of FL-S297A. Together, these data indicate that VbP stimulation does indeed disrupt the DPP9-binding checkpoint in cells.

Based on our structure and *in vitro* experiments, we posited that VbP-induced disruption is likely indirect and therefore reasoned it could only be observed in a cell-based system. To test this hypothesis, we ectopically expressed dTAG-CARD8-FL or dTAG-CARD8-ZUC with FLAG-tagged FIIND-SA in HEK 293T cells. The cells were then treated with dTAG-13 and/or VbP for 18 h prior to immunoprecipitation. dTAG-13 treatment induced degradation of dTAG-CARD8 and increased the amount of DPP9 captured by FIIND-SA-FLAG, likely due to increased CARD8-CT that may enhance FIIND-SA binding to DPP9 (Figure 5G). Notably, this effect was completely nullified by VbP (Figure 5G), even though VbP does not directly displace bound CARD8 to DPP9. We speculate that inhibition of DPP9/8 catalytic activity causes secondary effects on the DPP9-ternary complex in cells, as discussed below. Overall, these data reveal that VbP activates the CARD8 inflammasome by two signals, induction of N-terminal degradation and disruption of the DPP9-ternary complex (Figure 5H).

## DISCUSSION

In this study, we used a combination of structural, biochemical and cell biological methods to dissect the molecular basis of CARD8 regulation by DPP9. Notably, CARD8 and NLRP1, which are related in their FIIND-mediated autoprocessing, DPP9-mediated suppression, and DPP8/9 inhibitor-mediated activation, share important similarities and differences. Both NLRP1 and CARD8 use a full-length FIIND and DPP9 (at site A) to capture a free CT containing the UPA subdomain of the FIIND (at site B), thereby forming a ternary complex that suppresses CT oligomerization and activation. However, the NLRP1-CT inserts into the DPP9 active site, but the CARD8-CT does not. This difference in binding modes is intriguing, and may be explained by the N-terminal residues in NLRP1-CT, which possess a DPP9 substrate-like sequence (Ser-Pro); whereas the N-terminal residues in CARD8-CT are Ser-Leu (Figure 1A). Due to this structural difference, the NLRP1-CT, but not the CARD8-CT, is directly displaced by DPP9 inhibitors in vitro (at site B), clarifying previously observed results. Interestingly, while also having a Ser-Pro sequence at the N-terminus of its CT, mouse NLRP1 does not appear to be displaced by VbP (Griswold et al., 2019). Consistent with this observation, in the cryo-EM structure of the highly homologous rat NLRP1 in complex with DPP9, the N-terminal residues of the CT inserts into the DPP9 substrate channel but do not make specific interactions with the active site and the first few residues of the CT including the Ser-Pro sequence are disordered (Huang et al., 2020). Furthermore, despite lack of displacement of CARD8-CT from DPP9 by VbP *in vitro*, VbP treatment still reduced the association between DPP9 and CARD8-CT in cells to enhance CARD8 inflammasome activation, suggesting that DPP9 inhibitors nevertheless impact CARD8 binding. We speculate that the inhibited form of DPP9 may not “catch” free CT as well as the active form of DPP9, even if inhibitors cannot disrupt preformed complexes, or that DPP8/9 inhibition causes other cellular effects such as substrate accumulation to compete with CARD8 for DPP9 interaction. The mechanistic basis for this complex disruption in cells warrants further study.

In addition to these structural differences, CARD8 and NLRP1 exhibit distinct expression patterns, as noted above, and likely have different cellular functions. For example, CARD8 and NLRP1 may have evolved to respond to different pathogens. Consistent with this idea, recent reports revealed that human rhinovirus 3C protease and HIV protease directly cleave and activate human NLRP1 and CARD8, respectively (Robinson et al., 2020; Tsu et al., 2020; Wang et al., 2020). Moreover, human NLRP1 was also recently reported to sense intracellular long doublestranded RNAs by directly binding them through its leucine-rich repeat (LRR) domain (Bauernfried et al., 2020), which is absent in CARD8. However, it should be noted that our studies suggest that CARD8 and NLRP1 may have also both evolved to sense a single endogenous danger signal triggered by DPP8/9 inhibition. Another important difference between CARD8 and NLRP1 is the structure of assembled inflammasome: while CARD8 can only directly recruit caspase-1 and activate rapid pyroptosis without causing cytokine maturation, NLRP1 recruits caspase-1 via ASC to elicit an inflammatory signaling cascade with cytokine secretion and pyroptotic cell death (Ball et al., 2020; Hollingsworth et al., 2020a; Qin et al., 2020). In this context, it is possible that NLRP1 activation is more pro-inflammatory, as exemplified by the association of NLRP1 hyperactivation with a series of severe skin pro-inflammatory diseases (Grandemange et al., 2017; Jin et al., 2007; Levandowski et al., 2013; Zhong et al., 2016; Zhong et al., 2018). By contrast, CARD8 activation is more pro-death, shown by its ability to kill T cells and monocytic cancer cells without eliciting cytokine production (Adams et al., 2004; Johnson et al., 2020; Johnson et al., 2018; Linder et al., 2020; Okondo et al., 2016; Spagnuolo et al., 2013; Walsh et al., 2013).

In summary, diverse pathogenic signals and the cellular consequence of DPP8/9 inhibition induce the N-terminal degradation of NLRP1 and CARD8, but this does not necessarily result in inflammasome formation. Instead, the freed CT is captured by a full-length NLRP1 or CARD8 and DPP8/9, thereby preventing spurious inflammasome activation in cases with a low level of N-terminal degradation. It is tempting to speculate that this ternary complex might also offer a second point of regulation, as signals that strengthen this interaction would enhance CARD8 or NLRP1 inhibition, while signals that weaken this interaction (e.g., DPP8/9 inhibition) would promote CARD8 or NLRP1 activation. As CARD8 and NLRP1 interact differently the DPP9 active site channel, it is furthermore possible that these signals could impact CARD8 and NLRP1 in distinct ways. More mechanistic studies are required to further address this question.

## STAR✶METHODS

Detailed methods are provided in the online version of this paper and include the following:

- KEY RESOURCES TABLE
- CONTACT FOR REAGENT AND RESOURCE SHARING
- EXPERIMENTAL MODEL AND SUBJECT DETAILS
  - HEK 293T Cell Lines
  - Expi293F Cell Line
  - Sf9 Cell Line
  - THP-1 Cell Lines
- METHOD DETAILS
  - Constructs and Cloning
  - Cell Culture
  - Protein Expression and Purification
  - Cryo-EM Screening and Data Collection
  - Cryo-EM Data Processing
  - Atomic Model Building
  - Negative Stain Electron Microscopy
  - Immunoprecipitation Assays
  - LDH Cytotoxicity Assay
  - dTAG Assays
- QUANTIFICATION AND STATISTICAL ANALYSIS
- DATA AND SOFTWARE AVAILABILITY

## SUPPLEMENTAL INFORMATION

Supplemental information includes four figures and one table and can be found with this article online.

## ACKNOWLEDGEMENTS

We thank Dr. Maria Ericsson at the HMS EM facility for advice and training and Drs. Richard Walsh, Sarah Sterling, and Shaun Rawson at the Harvard Cryo-EM Center for Structural Biology for cryo-EM training and productive consultation. We thank Drs. Kangkang Song and Kyounghwan Lee for data collection at UMass Medical School. We thank Daija Bobe and Dr. Ed Eng for data collection at the National Center for Cryo-EM Access and Training (NCCAT) and the Simons Electron Microscopy Center located at the New York Structural Biology Center, supported by the NIH Common Fund Transformative High Resolution Cryo-Electron Microscopy program (U24 GM129539,) and by grants from the Simons Foundation (SF349247) and NY State. We thank Theo Humphreys and the Pacific Northwest Center for Cryo-EM (PNCC) at Oregon Health & Science University for screening and preliminary dataset collection, under the NIH grant U24GM129547 and accessed through EMSL (grid.436923.9), a DOE Office of Science User Facility sponsored by the Office of Biological and Environmental Research. We also thank Drs. Irina Novikova, Harry Scott, and Craig Yoshioka at PNCC for cryo-EM data processing training. pInducer20 was a gift from Stephen Elledge (Addgene plasmid # 44012). pLEX_307 was a gift from David Root (Addgene plasmid # 41392). pcDNA3.1 LIC 6A and pcDNA 3.1 LIC 6D were gifts from Scott Gradia (Addgene plasmid #30124 and #30127). pLEX_305-N-dTAG was a gift from James Bradner & Behnam Nabet (Addgene plasmid #91797) This work was supported by National Institutes of Health grants DP1 HD087988 and R01 AI124491 to H.W., T32-GM007726 to L.R.H., T32 GM007739-Andersen and F30 CA243444 to A.R.G., the Memorial Sloan Kettering Cancer Center Core Grant P30 CA008748 and R01 AI137168 to D.A.B. We thank BioRender for figure design and SBGrid for computing support.

## AUTHOR CONTRIBUTIONS

H.W., H.S., and L.R.H. conceived the project idea and designed the study. L.R.H. designed constructs with H.S.’ input. L.R.H. carried out preliminary expression and purification studies. L.R.H. purified the complexes. H.S. and L.R.H. made cryo-EM grids for data collection. L.R.H. screened grids and collected cryo-EM data. H.S. analyzed cryo-EM data with assistance from L.R.H. and performed model building and refinement for the DPP9-CARD8-WT complexes. L.R.H. analyzed cryo-EM data for the DPP9-CARD8-S297A complex. H.S. and L.R.H. designed mutants. L.R.H., and A.R.G. cloned mutants for *in vitro* and cell-based assays. A.R.G. performed all cell-based assays with assistance from J.C.H and Q.W. under D.A.B.’s supervision. A.R.G., L.R.H., H.S., H.W., and D.A.B. designed the experiments. L.R.H., A.R.G., D.A.B., and H.W. wrote the manuscript with input from all authors.

## STAR✶METHODS

### KEY RESOURCES TABLE

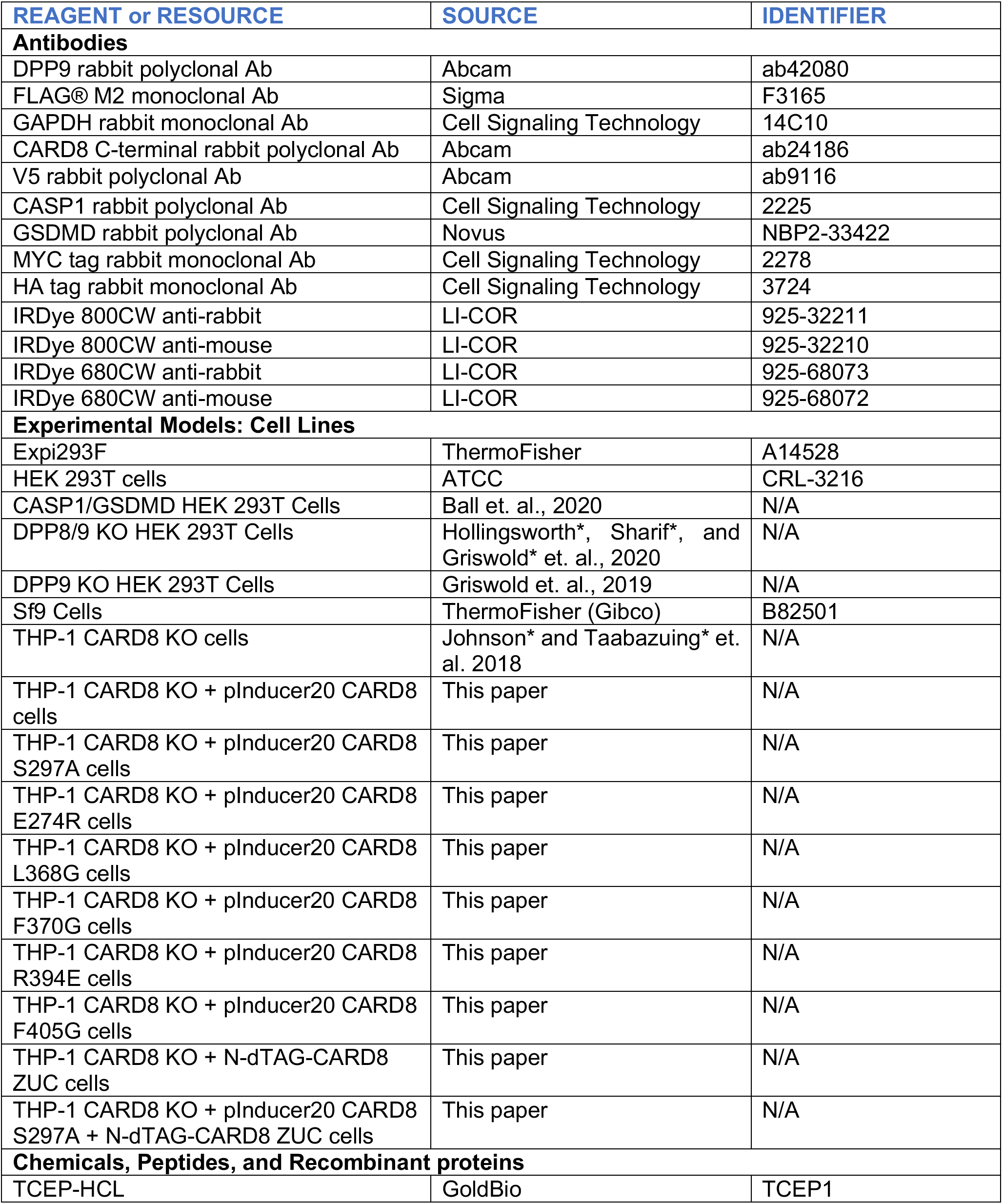
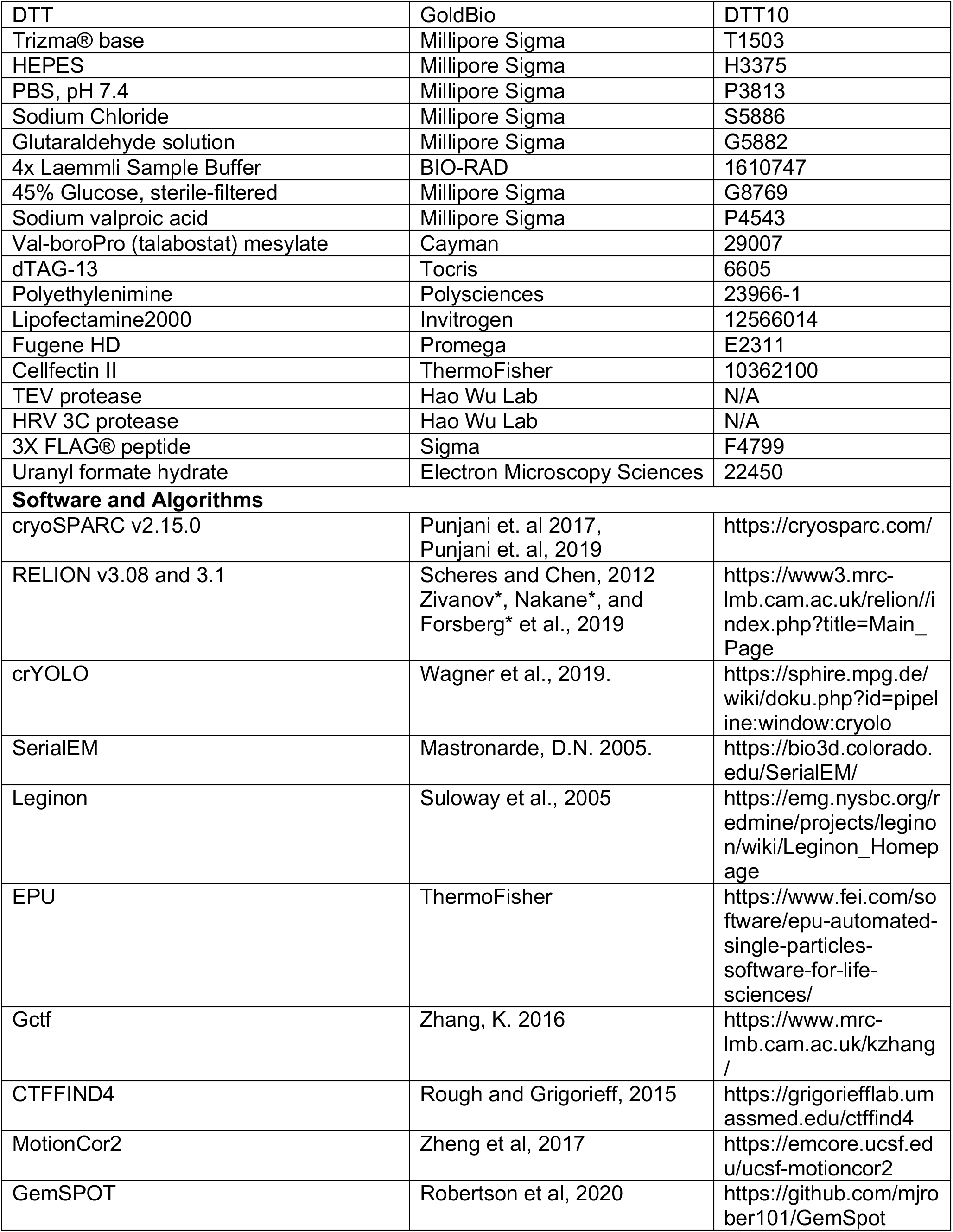
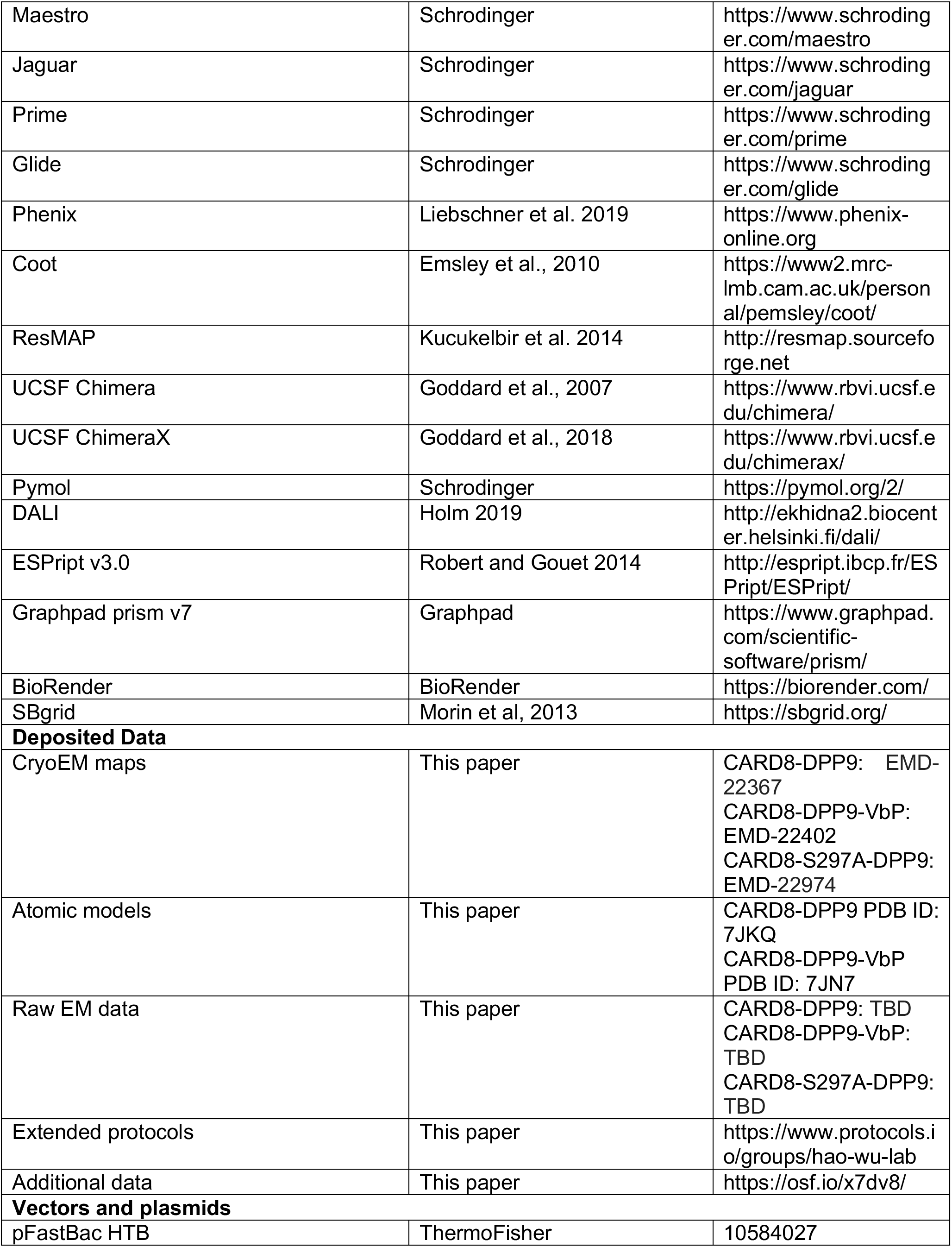
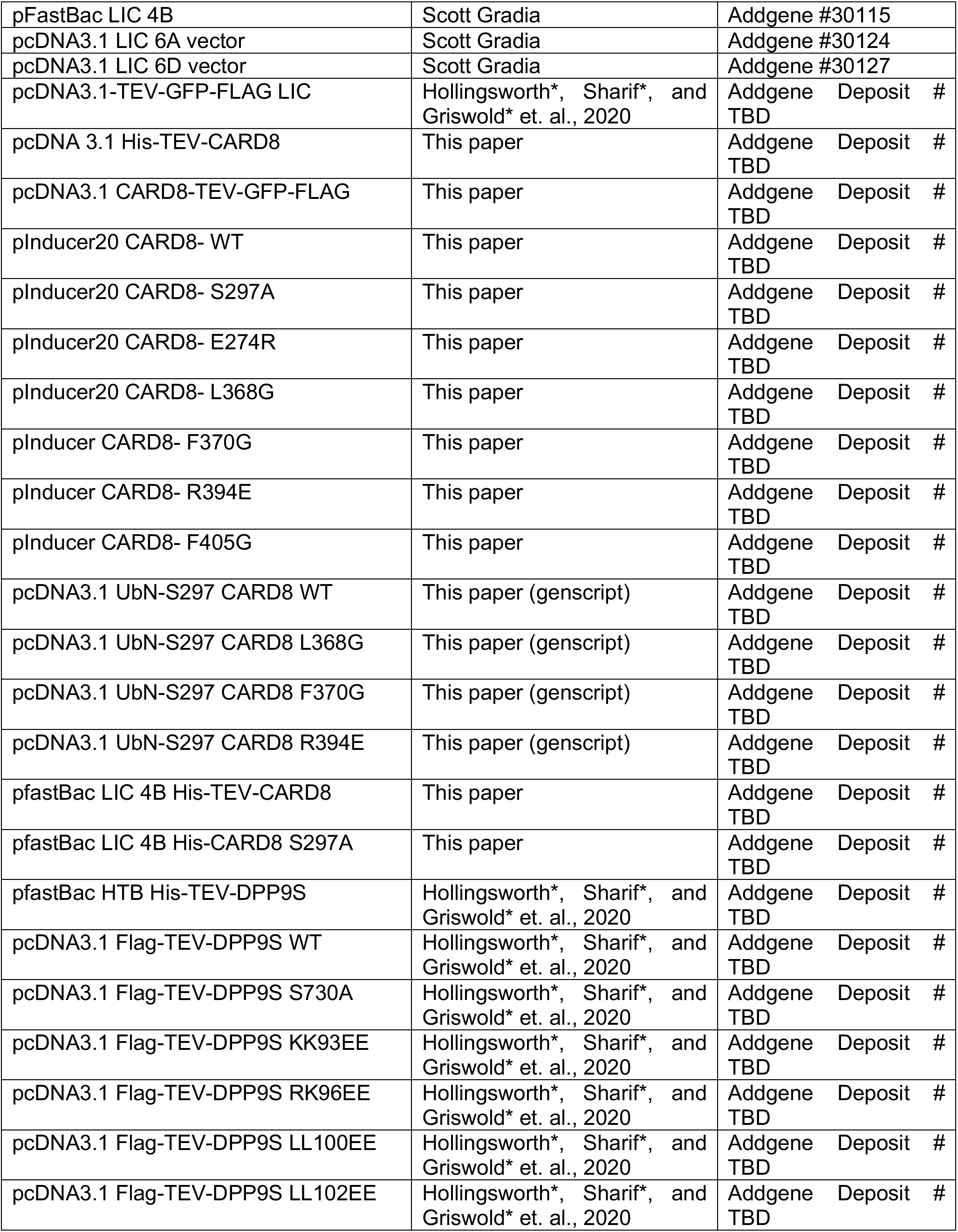
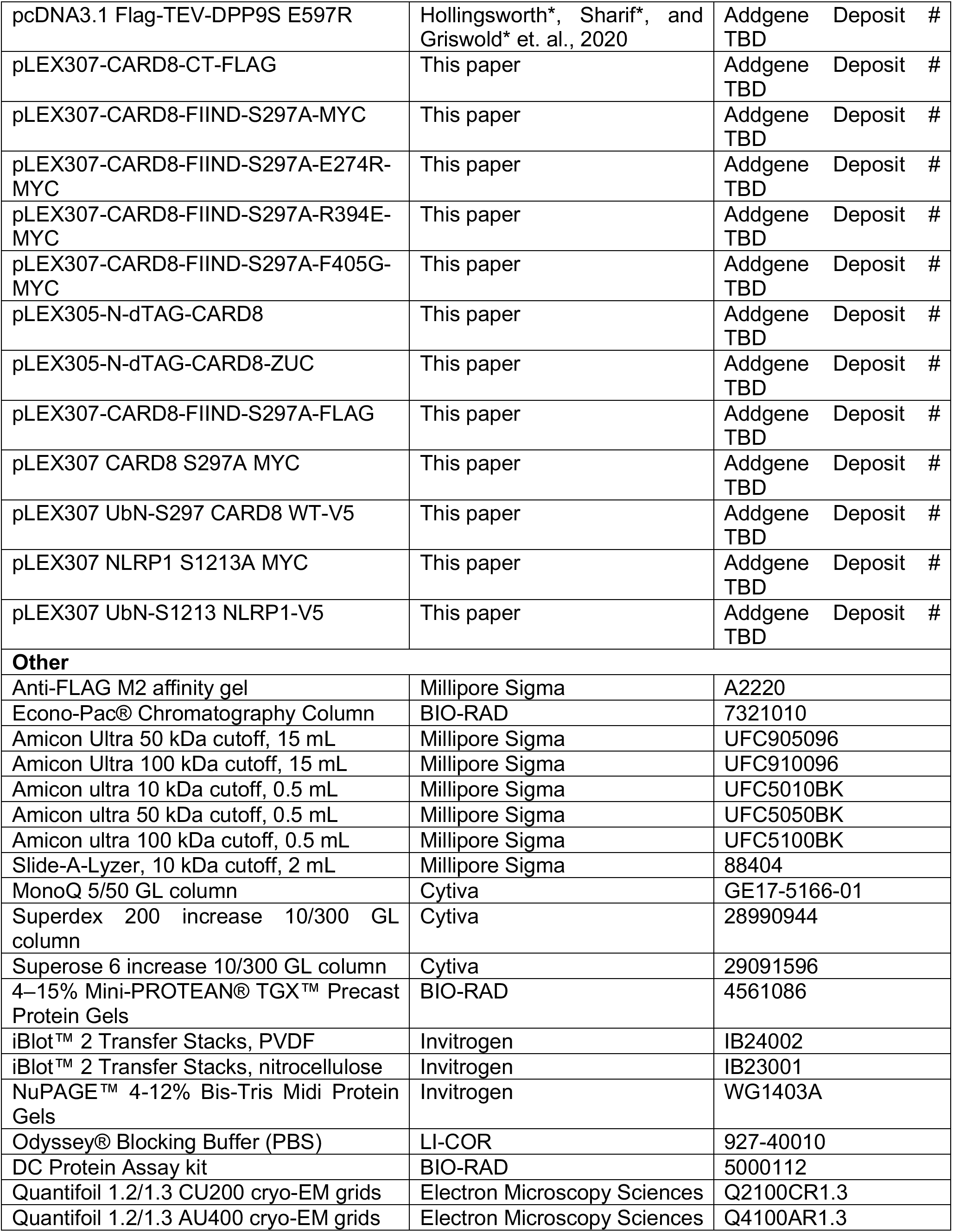
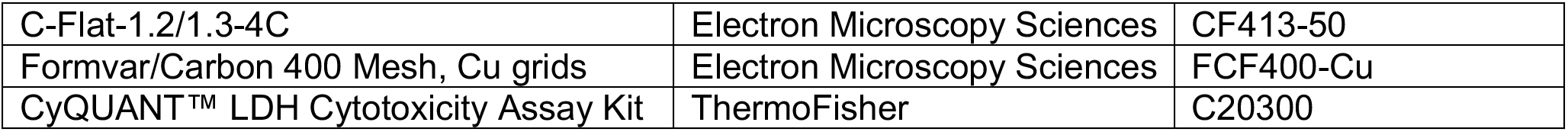

### CONTACT FOR REAGENT AND RESOURCE SHARING

Further information and requests for resources and reagents should be directed and will be fulfilled by the Lead Contact, Hao Wu. Study plasmids will be made available on Addgene. Extended protocol are available on protocols.io (https://www.protocols.io/groups/hao-wu-lab). Pymol and chimera session files, in addition to other raw data, will be made available on our Open Science Framework (https://osf.io/x7dv8/). Cryo-EM data were deposited to EMPIAR, the EMDB, and the PDB.

### EXPERIMENTAL MODEL AND SUBJECT DETAILS

#### HEK 293T Cell Lines

The human kidney epithelial cell line HEK 293T (ATCC) and related CASP1/GSDMD-expressing stable cell lines and DPP8/9 KO lines were maintained in DMEM (GIBCO, ThermoFisher) supplemented with 10% fetal bovine serum (GIBCO, ThermoFisher Scientific), at 37°C, and 5% CO2. HEK 293T cells were verified by the manufacturer. Cells were frequently checked for morphological features and tested for mycoplasma using MycoAlert Mycoplasma Detection kit (Lonza).

#### Expi293F Cell Line

Expi293F suspension cells were maintained in Expi293F Expression Medium (GIBCO, ThermoFisher) with constant shaking at 100 RPM, 37°C, 5% CO2. Expi293F cells were not authenticated nor tested for mycoplasma contamination.

#### Sf9 Cell Line

Sf9 insect cells were maintained in HyClone SFX-Insect Cell Media (Cytiva) supplemented with 1X antibiotic-antimycotic (ThermoFisher) at 27°C with constant shaking at 100 RPM. Sf9 cells were recently purchased from the manufacturer and were not authenticated.

#### THP-1 Cell Lines

THP-1 cells (ATCC) and related THP-1 CARD8^-/-^ stable cell lines were grown in Roswell Park Memorial Institute (RPMI) medium 1640 with L-glutamine and 10% fetal bovine serum (GIBCO, ThermoFisher Scientific), at 37°C and 5% CO2. Cells were monitored for morphological features and regularly tested for mycoplasma using MycoAlert Mycoplasma Detection kit (Lonza).

### METHODS DETAILS

#### Constructs and Cloning

Full-length CARD8 (T60 isoform, Uniprot ID Q9Y2G2-5) was cloned into pcDNA3.1 LIC 6A (Addgene plasmid #30124) with a C-terminal FLAG tag and a modified pcDNA3.1 LIC 6D (Addgene plasmid #30127) construct (C-terminal TEV-GFP-FLAG tag). The short isoform of DPP9 (DPP9S, Uniprot ID Q86TI2-1) was also cloned into pcDNA3.1 LIC 6A (N-terminal FLAG-TEV tag or N-terminal His-TEV tag). CARD8 was also shuttled into pInducer20 vector (Addgene, #44012) using Gateway technology (Thermo Fischer Scientific) and pLEX_305-N-dTAG (Addgene, #91797). CARD8-ZUC containing the ZU5, UPA and CARD was cloned as previously described (Chui et al., 2020) and shuttled to pLEX_305-N-dTAG (Addgene, #91797). CARD8-CT constructs were synthesized (GenScript) with an N-terminal ubiquitin sequence followed by CARD8 (S297-L537), cloned into the pcDNA3.1 vector (UbN-CARD8-CT) and shuttled into the pLEX307 vector using Gateway technology (Thermo Fischer Scientific). Point mutations were introduced with Q5 site-directed mutagenesis (NEB) or QuikChange site-directed mutagenesis (Agilent). All constructs will be made available on Addgene.

#### Cell Culture

HEK 293T cells and THP-1 cells were purchased from ATCC. HEK 293T cells were grown in Dulbecco’s Modified Eagle’s Medium (DMEM) with L-glutamine and 10% fetal bovine serum (FBS). THP-1 cells were grown in Roswell Park Memorial Institute (RPMI) medium 1640 with L-glutamine and 10% FBS. All cells were grown at 37° C in a 5% CO2 atmosphere incubator. Cell lines were regularly tested for mycoplasma using the MycoAlert Mycoplasma Detection Kit (Lonza). THP-1 CARD8^-/-^ cells and HEK 293T cells stably expressing caspase-1 and GSDMD were generated as previously described (Johnson et al., 2020). Reconstituted THP-1 CARD8^-/-^ cells were generated via lentiviral infection of THP-1 CARD8^-/-^ cells with the indicated pInducer20 construct followed by selection with G418 (Geneticin) at 200 μg/mL until all control cells were dead (approximately 14 days). dTAG-ZUC THP-1 cells were generated via lentiviral infection followed by selection with puromycin 500 ng/mL until all control cells were dead (approximately 7 days).

#### Protein Expression and Purification

To express CARD8-DPP9 complexes, Expi293F cells (1 L, 2-3 x 10^6^ cells/mL) were cotransfected with CARD8-TEV-GFP-FLAG (0.7 mg) and DPP9S (0.3 mg) following incubation with polyethylenimine (3 mL, 1 mg/mL) in OptiMEM (100 mL) for 30 min. 24 h later, cells were supplemented with glucose (9 mL, 45%) and valproic acid (10 mL, 300 mM). Cells were harvested 5 d after transfection by centrifugation (2,000 RPM, 20 min), washed once with PBS, split into 3 pellets, flash-frozen in liquid nitrogen, and stored at −80 °C. Later, one thawed pellet was resuspended in lysis buffer (50-100 mL, 25 mM Tris-HCl pH 7.5, 150 mM NaCl, 1 mM TCEP), sonicated (2 s on 8 s off, 3.5 min total on, 40% power, Branson), and ultracentrifuged at 40,000 RPM for 1 h (45 Ti fixed-angle rotor, Beckman). The supernatant was incubated with preequilibrated (lysis buffer) anti-FLAG M2 affinity gel (Sigma, 1.0 mL) for 4 h at 4 °C, washed in batch once with lysis buffer (5 mL), and then washed by gravity flow with 25-50 column volumes (CV) lysis buffer. The CARD8-DPP9 complex was eluted by on-column cleavage at room temperature for 1 h using elution buffer (5 mL, 25 mM Tris-HCl pH 7.5, 150 mM NaCl, 1 mM TCEP, 0.2 mg TEV protease) and loaded onto a Mono Q 5/50 GL anion exchange column (Cytiva). Protein was eluted using a salt gradient from 150 mM to 1 M NaCl (25 mM Tris-HCl pH 8.0, 1 mM TCEP) over 15 CV. Mono Q eluent was concentrated using a 0.5 mL spin concentrator (Amicon Ultra, 50 kDa MW cutoff) to 0.5 mg/mL (assuming ε=1). Concentrated eluent was dialyzed overnight into EM buffer (25 mM HEPES pH 7.5, 150 mM NaCl, 1 mM TCEP) using a 0.5 mL Slide-A-Lyzer (ThermoFisher). Total protein yield varied between 2-3 mg per L of mammalian culture. Expression and purification of CARD8 (S297A) followed an identical protocol. Expression and purification of the VbP-bound complex followed an identical protocol except for the addition of 10 μM VbP to all purification and dialysis buffers.

#### Cryo-EM Screening and Data Collection

Grids were screened at University of Massachusetts Worcester, Pacific Northwest Center for Cryo-EM (PNCC), and Harvard Medical school (HMS) using a Talos Arctica microscope (ThermoFisher) operating at an acceleration voltage of 200 keV equipped with a direct electron detector. Small initial dataset collection revealed severe particle orientation preference for the CARD8-DPP9 maps, causing anisotropic resolution, in addition to dissociation of CARD8-DPP9 complexes. To remedy these issues, we crosslinked samples to avoid dissociation and tilted the stage to collect missing views (below).

The purified DPP9S-CARD8-WT or DPP9S-CARD8-S297A complex (0.40 mg/mL assuming ε=1; 25 mM HEPES pH 7.5, 150 mM NaCl, 1 mM TCEP, ±10 μM VbP) was crosslinked with 0.02% glutaraldehyde on ice for 5 min and immediately loaded onto a glow-discharged Quantifoil grid (R1.2/1.3 400-mesh gold-supported holey carbon, Electron Microscopy Sciences), blotted for 3–5 s under 100% humidity at 4 °C, and plunged into liquid ethane using a Mark IV Vitrobot (ThermoFisher). Grids were screened at Harvard Medical School for ice and particle quality prior to data collection. For data collection, movies were acquired at Harvard Medical School (DPP9-CARD8-WT) and National Center for CryoEM Access and Training (NCCAT) in New York (DPP9-CARD8-WT+VbP and DPP9-CARD8-S297A) using a Titan Krios microscope (ThermoFisher) at an acceleration voltage of 300 keV equipped with a BioQuantum K3 Imaging Filter (slit width 20 eV). Movies were recorded with a K3 Summit direct electron detector (Gatan) operating in counting mode at 105,000 x (0.825 Å/pix at HMS, 0.82 or 0.83 at Å/pix NCCAT).

For CARD8-DPP9 at 0° and 37°: 3,306 and 2,488 movies at a stage tilt of either 0° or 37° were collected using SerialEM (Mastronarde, 2005) at varying defocus levels ranging between −0.8 to −2.2 μm and −1.5 to −3.0 μm, respectively. We used image shift to record two shots for each of the four holes per stage movement. Movies were exposed with a total dose of 58.5 e^-^/Å^2^ for 2.22 s fractionated over 49 frames for 0° stage tilt movies and 64.99 e^-^/Å^2^ for 2.25 s fractionated over 49 frames for 37° stage tilt movies.

For CARD8-DPP9-VbP at 0° and 37°: 1,811 and 6,642 movies at a stage tilt of either 0° or 37°, respectively, were collected using Leginon (Suloway et al., 2005) at varying defocus levels ranging between −1.1 to −3.4 μm. One shot was recorded for each of the four holes per stage movement through image shift. All movies were exposed with a total dose of 67.06 e^-^/Å^2^ for 1.5 s fractionated over 50 frames.

For CARD8(S297A)-DPP9, 3,840 movies were collected at a stage tilt of 37° using Leginon (Suloway et al., 2005) to vary the defocus range between −0.8 to −2.5 μm and to record one shot for each of the four holes per stage movement through image shift. All movies were exposed with a total dose of 63.67 e^-^ /Å^2^ for 1.5 s fractionated over 50 frames.

#### Cryo-EM Data Processing

Data processing leveraged SBgrid Consortium (Morin et al., 2013) for support and computing resources. Movies collected at Harvard Medical School (CARD8-DPP9) were pre-processed on-the-fly by the facility’s pipeline script. Movies were corrected by gain reference and for beam-induced motion and summed into motion-corrected images using the Relion 3.08 implementation of the MotionCor2 algorithm (Zheng et al., 2017). The CTFFIND4 program (Rohou and Grigorieff, 2015) was used to determine the defocus of each micrograph. Relion 3.1 (Scheres, 2012; Zivanov et al., 2018) was used for subsequent image processing. Movies collected at NCCAT (CARD8-DPP9-VbP) were similarly pre-processed using Relion 3.1.

For the CARD8-DPP9 complex, template-free autopicking with crYOLO (Wagner et al., 2019) (generalized training for on-the-fly picking at the HMS cryo-EM center) selected 487,952 particles from 3,306 micrographs, which were subjected to a single round of 2D classification. The heterogenous nature of the sample was evident in 2D class averages because complexes containing only DPP9, 2CARD8:2DPP9, and 4CARD8:2DPP9 were present. A randomized set of 100,000 particles was selected for the *de novo* reconstruction of an initial model, which was low-pass-filtered to 40 Å to use as the input reference for 3D classification. Multiple rounds of 3D classification including global and local fine angular search were performed. After visual inspection, one class with 2CARD8:2DPP9 (62,018 particles) was selected for 3D refinement. Subsequently, particles were CTF refined and Bayesian polished to reach an overall resolution of 3.3 Å. However, the map suffered from anisotropic resolution, and cryoEF (Naydenova and Russo, 2017) analysis estimated that data collected at a tilt angle of 37° was ideal to fill the gaps in Fourier space.

The overall processing scheme for tilt data was derived from the work flow of tilt dataset described in Zivanov *et al.* 2018 (Zivanov et al., 2018). For CARD8-DPP9 data collected at a 37° tilt, micrographs were first motion-corrected with MotionCor2 (Zheng et al., 2017) followed by Gctf (Zhang, 2016) to calculate per micrograph defocus values in Relion 3.1. Template-free autopicking with crYOLO (Wagner et al., 2019) then picked 313,425 particles from 2,488 micrographs. After multiple rounds of 2D classification, 243,459 visually homogenous particles remained for further processing. A random subset comprising 121,730 of these particles was used for *de novo* initial model construction, which was then low-passed filtered (30 Å) and used as the initial reference map for 3D classification. After one round of 3D classification, a 3D class with 58,980 particles was selected for high-resolution refinement. The first 3D refinement using these particles yielded a 5.2 Å resolution structure and the Fourier shell correlation (FSC) curve showed strong fluctuations indicating imprecisions in CTF estimation. We then utilized CTF refinement implemented in Relion 3.1, including higher order aberrations correction, anisotropic magnification corrections, and per-particle defocus estimation (Zivanov et al., 2020). Iterative rounds of 3D refinement followed by CTF refinement and Bayesian polishing gradually improved the resolution and this iterative process was stopped when we observed a resolution plateau at 3.8 Å. Refined and polished particle sets from 0° and 37° stage-tilt data were merged and 3D refinement was performed. The merged data was then reconstructed to give an overall resolution of 3.3 Å as calculated by gold-standard FSC between half maps, with much improved CARD8 density (Figure S1).

For the CARD8-DPP9-VbP complex, template-free autopicking with crYOLO (Wagner et al., 2019) (trained with 10 manually-picked high-contrast micrographs) yielded 1,404,573 particles from 1,811 micrographs that were collected without stage tilt. These particles were subjected to multiple rounds of 2D classification that yielded visually homogeneous 2D classes with 471,255 particles. A randomized set of 100,000 particles was selected for the *de novo* reconstruction of an initial model, which was low-pass-filtered to 30 Å to use as the input reference for 3D classification. Multiple rounds of 3D classification including global and local fine angular search were performed. After visual inspection, one 3D class (89,909 particles) was selected for 3D refinement. Subsequently, particles were CTF refined and Bayesian polished to reach an overall resolution of 3.5 Å. However, the map suffered from anisotropic resolution, and cryoEF (Naydenova and Russo, 2017) analysis estimated that data collected at a tilt angle of 37° degrees was ideal to fill the gaps in Fourier space (below).

The overall processing scheme for tilt data analysis of CARD8-DPP9-VbP was derived from CARD8-DPP9 data processing (described above). Briefly, micrographs were first motion-corrected with MotionCor2 (Zheng et al., 2017) followed by Gctf (Zhang, 2016) to calculate per micrograph defocus values in Relion 3.1. Template-free autopicking with crYOLO (Wagner et al., 2019) then picked 1,889,993 particles from 6642 micrographs. After multiple rounds of 2D classification, 840,902 visually homogenous particles remained for further processing. A random subset comprising 100,000 of these particles was used for *de novo* initial model construction, which was then low-passed filtered (30 Å) and used as the initial reference map for 3D classification. After multiple rounds of 3D classification, a 3D class with 91,254 particles was selected for high-resolution refinement. The first 3D refinement using these particles yielded a 5.8 Å resolution structure, and the FSC curve showed strong fluctuations indicating imprecisions in CTF estimation. We then utilized CTF refinement implemented in Relion 3.1, including correction for higher order aberrations and anisotropic magnification, and per-particle defocus estimation (Zivanov et al., 2020). Iterative rounds of 3D refinement followed by CTF refinement and Bayesian polishing gradually improved the resolution and this iterative process was stopped when we observed a resolution plateau at 3.4 Å. Refined particle sets from 0° and 37° stage-tilt data were merged and 3D refinement was performed. The merged data was then 3D classified with fine local search angles to give a final stack of 146,101 particles which was reconstructed to give an overall resolution of 3.3 Å as calculated by gold-standard FSC between half maps, with much improved CARD8 density (Figure S4).

The DPP9-CARD8-S297A complex was primarily processed in cryoSPARC (Punjani et al., 2017). 3,840 movies collected at 37° stage tilt were summed into motion-corrected micrographs with Patch-Motion in cryoSPARC (Punjani et al., 2017). Next, defocus values were determined with cryoSPARC’s Patch-CTF function (Punjani et al., 2017). 5,505,288 particles were picked by blob picking and the 2D classes following two rounds of classification were used as input to repick 1,634,615 particles. Following 3 rounds of 2D classification, 191,609 particles remained and were used to generate 3 ab-initio models. At this stage, heterogeneity between bound and unbound DPP9 particles was evident, and a round of heterogeneous refinement (2 classes) yielded a reconstruction from 92,404 particles at a nominal resolution of 4.88 Å. These particles were used for a final round of non-uniform refinement (Punjani et al., 2019) which converged at an overall resolution of 3.86 Å as calculated by gold-standard FSC (Figure S3).

#### Atomic Model Building

The cryo-EM maps were first fit with the crystal structure of DPP9 dimer (PDB ID: 6EOQ) (Ross et al., 2018). A homology model of CARD8-FIIND was generated with Schrodinger Prime (Jacobson et al., 2002) using the structure of NLRP1-FIIND (Hollingsworth et al., 2020a; Qin et al., 2020) as template. Manual adjustment and *de novo* building of missing segments, rigid-body fitting, flexible fitting, and segment-based real-space refinement were performed in distinct parts of the initial model to fit in the density in Coot (Emsley et al., 2010), with help of UCSF-Chimera (Goddard et al., 2007) and real-space refinement in Phenix (Klaholz, 2019). A few unstructured regions, including parts of the UPA, were omitted owing to poor density. The full model represents Asp18-Met1356 amino acids of DPP9 (short isoform, Uniprot ID Q86TI2-1), ZU5^A^ Gly166-Ser295, UPA^A^ Arg304-Pro446 and UPA^B^ Leu320-Pro446 of CARD8 (isoform 5, Uniprot ID Q9Y2G2-5).

For the VbP-bound structure, the full model represents Asp18-Leu863 amino acids of DPP9 (short isoform, Uniprot ID Q86TI2-1), ZU5^A^ Gly166-Ser295, UPA^A^ Arg304-Pro445 and UPA^B^ Leu320-Pro446 of CARD8 (isoform 5, Uniprot ID Q9Y2G2-5). We modelled covalently linked DPP9 S730-VbP in the cryo-EM map density using NLRP1-DPP9-VbP (PDB ID: 6X6C) and DPP8-VbP (PDB ID: 6HP8) (Díaz, 2018) as templates. DPP9-VbP interactions are similar to those observed in the DPP8-VbP complex in addition to DPPIV bound to substrate (PDB ID: 5YP3) (Roppongi et al., 2018) (Figure S4G).

For both structures, interaction analysis was conducted visually and using PISA (Krissinel and Henrick, 2007). Structure representations were generated in ChimeraX (Goddard et al., 2018), Pymol, and ResMap (Kucukelbir et al., 2014). Ligand interaction analysis was conducted with Maestro (Schrödinger Release 2020-1, 2020). Pymol and ChimeraX session files are available on our Open Science Framework repository (https://osf.io/x7dv8/). Schematics were created with BioRender.

#### Negative Stain Electron Microscopy

Copper grids coated with layers of plastic and thin carbon film (Electron Microscopy Sciences) were glow discharged before 4 μl of purified proteins were applied. Samples were left on the grids for 45 sec, blotted, and then stained with 1% uranyl formate for 40 s, blotted, and air dried. The grids were imaged on a JEOL 1200EX or Tecnai G^2^ Spirit BioTWIN microscope at the HMS EM facility operating at 80 keV.

#### Immunoprecipitation Assays

For DPP9 mutants (Figure 3B), *DPP8/9 DKO* HEK 293T cells (Hollingsworth et al., 2020b) were seeded at 1 x 10^6^ cells/well in 6-well tissue culture dishes. The following day cells were transfected with plasmids encoding for CARD8 (2 μg) or the indicated FLAG-tagged DPP9 construct (2 μg) with FuGENE HD according to manufacturer’s instructions (Promega). Cells were harvested and washed 3x with PBS. Pellets were lysed in Tris-Buffered Saline (TBS) with 0.5% NP-40 using pulse sonication and centrifuged at 20,000 x *g* for 10 min at 4 °C. DPP9 and CARD8 lysates were mixed in a 1:1 ratio prior to treating with DMSO or VbP (10 μM) for 1 h. They were incubated with 20 μL of anti-FLAG-M2 agarose resin (Sigma) overnight at 4 °C. After washing 3 x 500 μL with cold PBS in microcentrifuge spin columns (Pierce), bound proteins were eluted by incubating resin with 40 μL of PBS with 150 ng/μL 3x-FLAG peptide for 1 h at 4 °C. An equal volume of 2x sample loading was added to the eluate and boiled. Protein content was evaluated by immunoblotting with the following antibodies: DPP9 rabbit polyclonal Ab (Abcam, Ab42080), FLAG^®^ M2 monoclonal Ab (Sigma, F3165), CARD8 C terminus rabbit polyclonal Ab (Abcam, ab24186), and GAPDH rabbit polyclonal Ab (CST, 14C10). Immunoblots were developed with the Odyssey CLx imaging system (LI-COR).

For CARD8 mutants (Figure 3C), THP-1 CARD8^-/-^ cells stably expressing the indicated pInducer20 CARD8 construct (20 x 10^6^ cells in 10 mL) were treated with doxycycline (1 μg/mL) and the pan-caspase inhibitor Z-VAD-FMK (20 μM) for 24 h. Cells were harvested and washed 3x with PBS. Pellets were lysed in Tris-Buffered Saline (TBS) with 0.5% NP-40 using pulse sonication and centrifuged at 20,000 x *g* for 10 min at 4 °C. Lysates were incubated with 50 μL of anti-HA agarose resin (Thermo Fisher Scientific) for 4 h at 4 °C. After washing 3 x 500 μL with cold PBS, bound proteins were eluted by boiling in 100 μL of 2x sample loading dye. Protein content was evaluated by immunoblotting with the following antibodies: DPP9 rabbit polyclonal Ab (Abcam, Ab42080), HA-Tag (C29F4) rabbit monoclonal (CST, #3724) and GAPDH rabbit polyclonal Ab (CST, 14C10).

For ternary complex capture experiments (Figure 4A), HEK 293T cells were seeded at 5 x 10^5^ cells/well in 6-well tissue culture dishes. The following day the cells were transfected with plasmids encoding for FLAG-tagged CARD8-CT (1 μg), the indicated MYC-tagged CARD8-FIIND-SA (1 μg), and RFP (to 2 μg) with FuGENE HD according to manufacturer’s instructions (Promega). After 48 h cells were harvested and washed 3x with PBS. Pellets were lysed in Tris-Buffered Saline (TBS) with 0.5% NP-40 using pulse sonication and centrifuged at 20,000 x *g* for 10 min at 4 °C. Lysates were incubated with 20 μL of anti-FLAG-M2 agarose resin (Sigma) overnight at 4 °C. After washing 3 x 500 μL with cold PBS in microcentrifuge spin columns (Pierce), bound proteins were eluted by incubating resin with 40 μL of PBS with 150 ng/μL 3x-FLAG peptide for 1 h at 4 °C. An equal volume of 2x sample loading was added to the eluate and boiled. Protein content was evaluated by immunoblotting with the following antibodies: DPP9 rabbit polyclonal Ab (Abcam, Ab42080), FLAG^®^ M2 monoclonal Ab (Sigma, F3165), Myc-tag (71D10) rabbit monoclonal Ab (CST, #2278), and GAPDH rabbit polyclonal Ab (CST, 14C10).

For on-bead displacement experiments (Figure 5A-C), HEK 293T cells were seeded at 3 x 10^6^ cells in a 10 cm tissue culture dish. The following day the cells were transfected with plasmids encoding for FLAG-tagged DPP9 (2 μg), V5-tagged UbN-S297 CARD8 or V5-tagged UbN-S1213 NLRP1 (3 μg), and MYC-tagged CARD8 S297A or MYC-tagged NLRP1 S1213A (5 μg) with FuGENE HD according to manufacturer’s instructions (Promega). Cells were harvested and washed 3x with PBS. Pellets were lysed in Tris-Buffered Saline (TBS) with 0.5% NP-40 using pulse sonication and centrifuged at 20,000 x *g* for 10 min at 4 °C. Lysates were incubated with 100 μL of anti-FLAG-M2 agarose resin (Sigma) for 2 h at 4 °C. The agarose was washed once with PBS, and subsequently split into 4 x 25 μL aliquots. 50 μL of PBS containing DMSO, VbP (10 μM), 8J (50 μM), Bestatin methyl ester (MeBS, 10 μM) at RT for 1 h in microcentrifuge spin columns (Pierce). Displaced proteins were collected via centrifugation. The resin was washed 3x with cold PBS, followed by elution with 3x-FLAG peptide as above. Protein content was evaluated by immunoblotting with the following antibodies: FLAG^®^ M2 monoclonal Ab (Sigma, F3165), Myc-tag (71D10) rabbit monoclonal Ab (CST, #2278), and V5 rabbit polyclonal Ab (Abcam, ab9116).

For dTAG ternary complex experiments (Figure 5G), HEK 293T cells were seeded at 5 x 10^5^ cells/well in 6-well tissue culture dishes. The following day the cells were transfected with plasmids encoding for FLAG-tagged CARD8-FIIND-S297A (1 μg), and dTAG-CARD8 (1 μg). The next day cells were treated with dTAG-13 (500 nM) and VbP (10 μM) for 24 h prior to harvesting. Anti-FLAG immunoprecipitation was executed as described above. Protein content was evaluated by immunoblotting with the following antibodies: FLAG^®^ M2 monoclonal Ab (Sigma, F3165), HA-Tag (C29F4) rabbit monoclonal (CST, #3724), DPP9 rabbit polyclonal Ab (Abcam, Ab42080), and GAPDH rabbit polyclonal Ab (CST, 14C10).

#### LDH Cytotoxicity Assay

For THP-1 CARD8^-/-^ cells stably expressing the indicated pInducer20 CARD8 construct (Figure 3D), cells were seeded at 2.5 x 10^5^ cells/mL in 12-well tissue culture dishes. The cells were treated with the indicated combination of doxycycline (1 μg/mL) and VbP (10 μg/mL) for 24 h. For HEK 293T cells stably expressing CASP-1 and GSDMD-V5 (Figure 3E), cells were seeded at 1.5 x 10^5^ cells/mL in 12-well tissue culture dishes. The following day the cells were transfected with plasmids encoding for the indicated CARD8-CT construct (0.1 μg) and RFP as a filler vector (to 2 μg) per 125 μL of Optimem with FuGENE HD according to manufacturer’s instructions (Promega).

In all cases, supernatants were analyzed for LDH activity using the CyQUANT™ LDH Cytotoxicity Assay Kit (ThermoFisher) and lysates protein content was evaluated by immunoblotting with a combination of the following antibodies: CARD8 CT rabbit polyclonal Ab (Abcam, Ab24186), CASP1 rabbit polyclonal Ab (CST, #2225), GSDMDC1 rabbit polyclonal Ab (Novus, NBP2-33422), HA-Tag (C29F4) rabbit monoclonal (CST, #3724) and GAPDH rabbit polyclonal Ab (CST, 14C10). Immunoblots were developed with the Odyssey CLx imaging system (LI-COR).

#### dTAG Assays

HEK 293T cells stably expressing CASP-1 and GSDMD-V5 (Figure 4C, E-F and Figure 5D-E) were seeded at 1.5 x 10^5^ cells/mL in 12-well tissue culture dishes. The following day the cells were transfected with plasmids encoding for the indicated amount of plasmids encoding for dTAG-CARD8 or dTAG-CARD8 ZUC and CARD8 FIIND-S297A constructs with RFP as a filler vector (to 2 μg) per 125 μL of Optimem with FuGENE HD according to manufacturer’s instructions (Promega). After 24 h (dTAG-CARD8) or 48 h (dTAG-CARD8 ZUC) cells were treated with DMSO, dTAG-13 (500 nM), and/or VbP (10 μM) for 3h prior to harvesting for analysis of supernatants for LDH release and lysates for protein content as detailed above.

THP-1 CARD8^-/-^ cells stably expressing dTAG-CARD8 ZUC (Figure 5F) were seeded at 2 x 10^5^ cells/mL in 12-well tissue culture dishes. Cells were treated with doxycycline (1 μg/mL) for 48h followed by dTAG-13 (5 nM) and VbP (10 μM) for 3h prior to harvesting for analysis of supernatants for LDH release as detailed above.

### QUANTIFICATION AND STATISTICAL ANALYSIS

Statistical significance was calculated by using GraphPad Prism version 7, GraphPad Software, San Diego, California USA, www.graphpad.com. The number of independent experiments, the statistical significance, and the statistical test used to determine the significance are indicated in each figure or figure legend or method section where quantification is reported.

### DATA AVAILABILITY

The cryo-EM maps have been deposited in the Electron Microscopy Data Bank under the accession numbers EMD-22367 (CARD8-DPP9), EMD-22402 (CARD8-DPP9-VbP), and EMD-22974 (CARD8-S297A-DPP9). The atomic coordinates have been deposited in the Protein Data Bank under the accession numbers PDB ID 7JKQ (CARD8-DPP9) and PDB ID 7JN7 (CARD8-DPP9-VbP). Raw cryo-EM data will be deposited into EMPIAR. Extended protein purification protocols will be made available on Protocols.io (https://www.protocols.io/groups/hao-wu-lab). Pymol session files and full-sized figures will be made available on OSF (https://osf.io/x7dv8/). All constructs will be made available on Addgene. All other data and materials can be obtained from the corresponding authors upon reasonable request.

## Supplemental Figures

**Figure S1.**
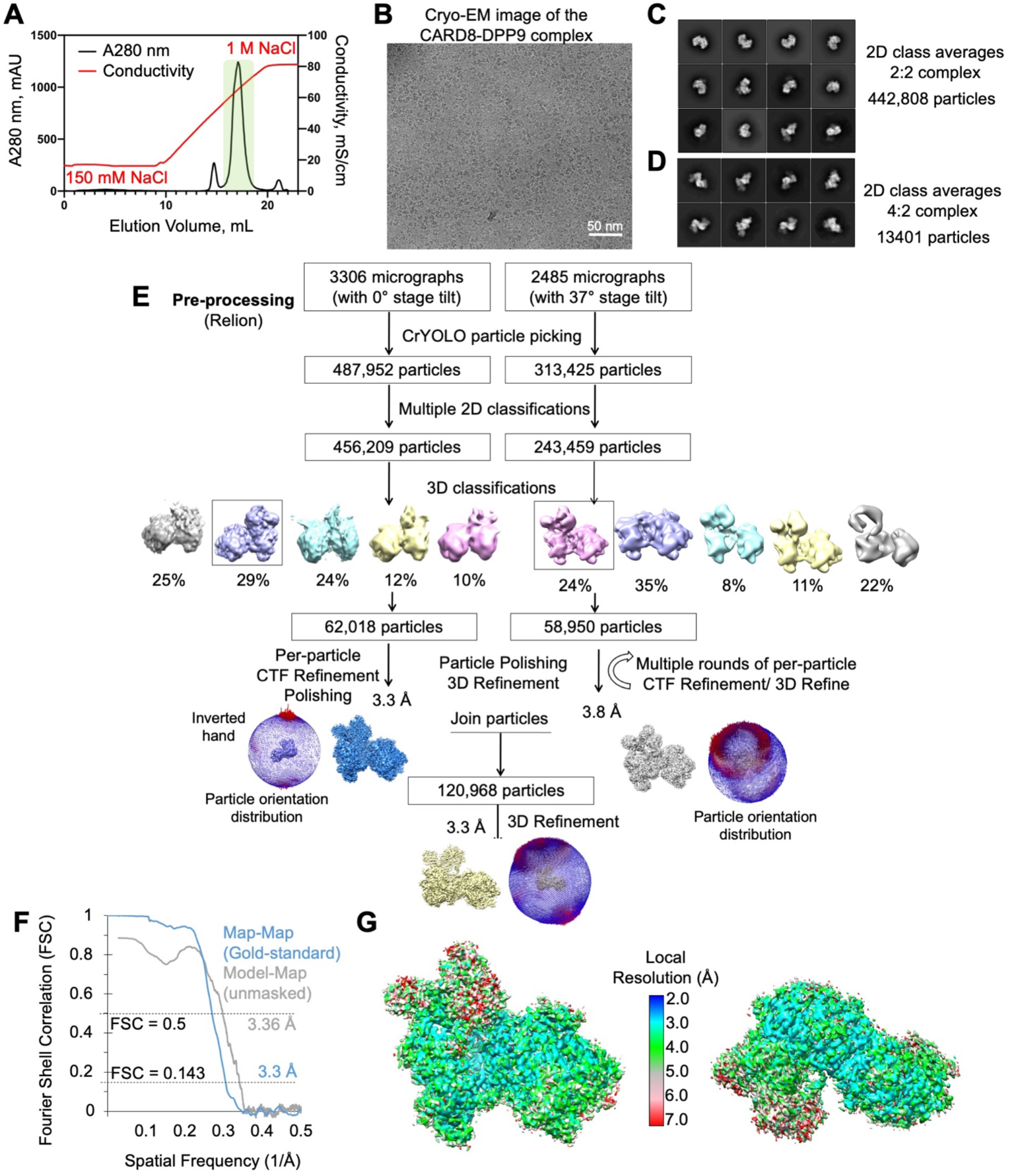
Cryo-EM Structure Determination of the DPP9-CARD8(WT) Complex, Related to Figure 1. (A) MonoQ anion exchange elution profile of the CARD8-DPP9 complex. Highlighted (green) peak indicates the fractions used for subsequent dialysis into lower salt (150 mM NaCl) buffer and cryoEM structure determination. (B) A representative cryo-EM micrograph. (C) Representative 2D class averages containing 2:2 CARD8:DPP9 complexes (D) Representative 2D class averages containing 4:2 CARD8:DPP9 complexes. (E) Workflow of cryo-EM data analysis with CrYOLO and RELION v3.1. Datasets collected at 0° and 37° stage tilts were merged to yield a cryo-EM map at 3.3 Å. (F) Gold-standard Fourier Shell Correlation (FSC) curves between the two independent half maps (blue) and between the unmask map and the atomic model (gray). (G) Local resolution of the cryo-EM map determined by ResMap and colored as indicated.

**Figure S2.**
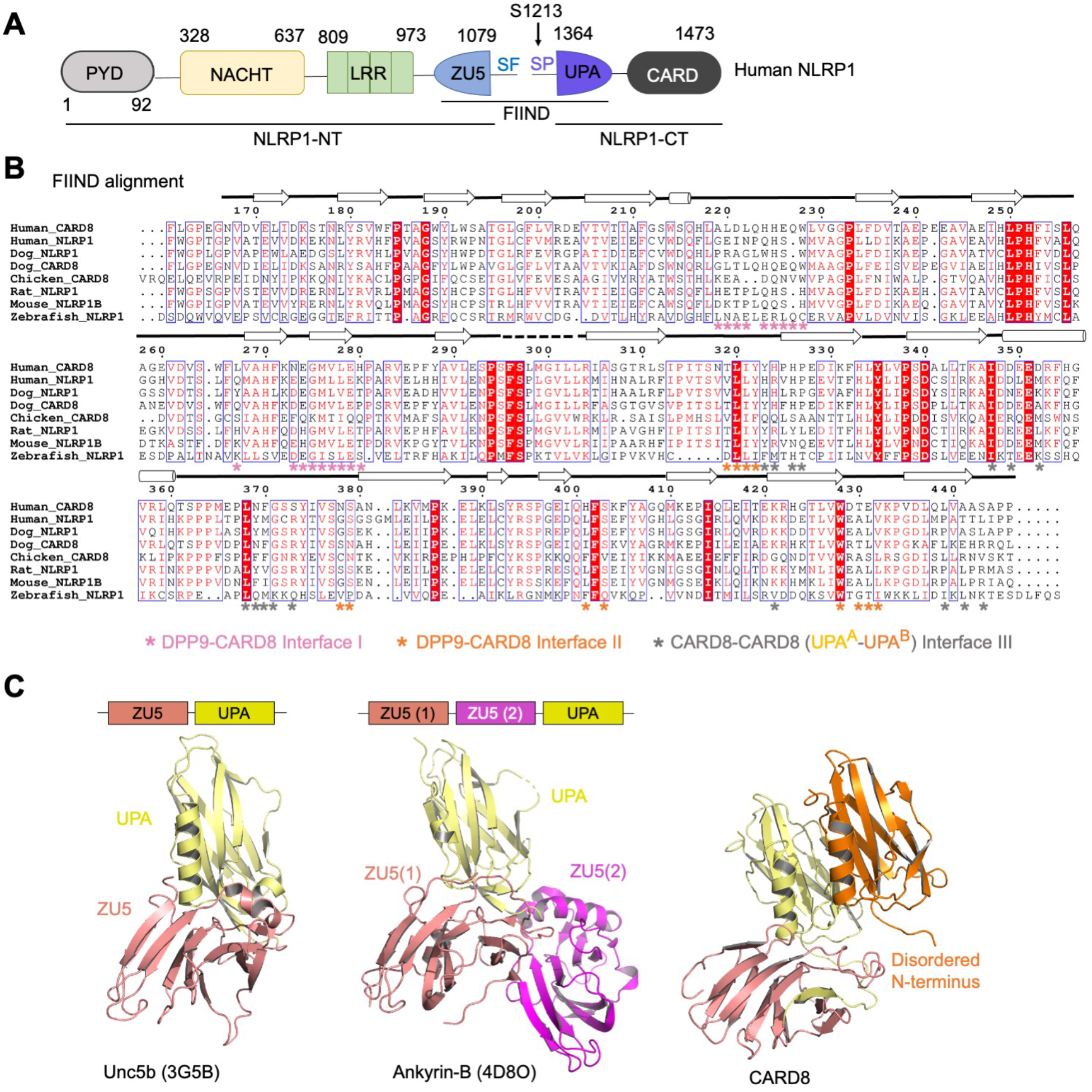
Comparison of ZU5 and UPA Folds Among Different Species and Protein Homologs, Related to Figure 1. (A) Domain organization of human NLRP1, highlighting the FIIND shared with CARD8. (B) Sequence alignment in the FIIND among CARD8 and NLRP1 homologs ClustalW alignment (Larkin et al., 2007). Chicken does not have NLRP1 annotated, whereas zebrafish, mouse, and rat are missing CARD8. Highly conserved residues are annotated by Esprint 3 (Robert and Gouet, 2014). Secondary structure and interfaces on the CARD8-DPP9 binding interface are annotated. (C)The ZU5-UPA domains from Unc5b (left) and the ZU5(1)-ZU5(2)-UPA domains from Ankyrin-B (right). CARD8 mimics the arrangement of ZU5(1)-UPA (right most).

**Figure S3.**
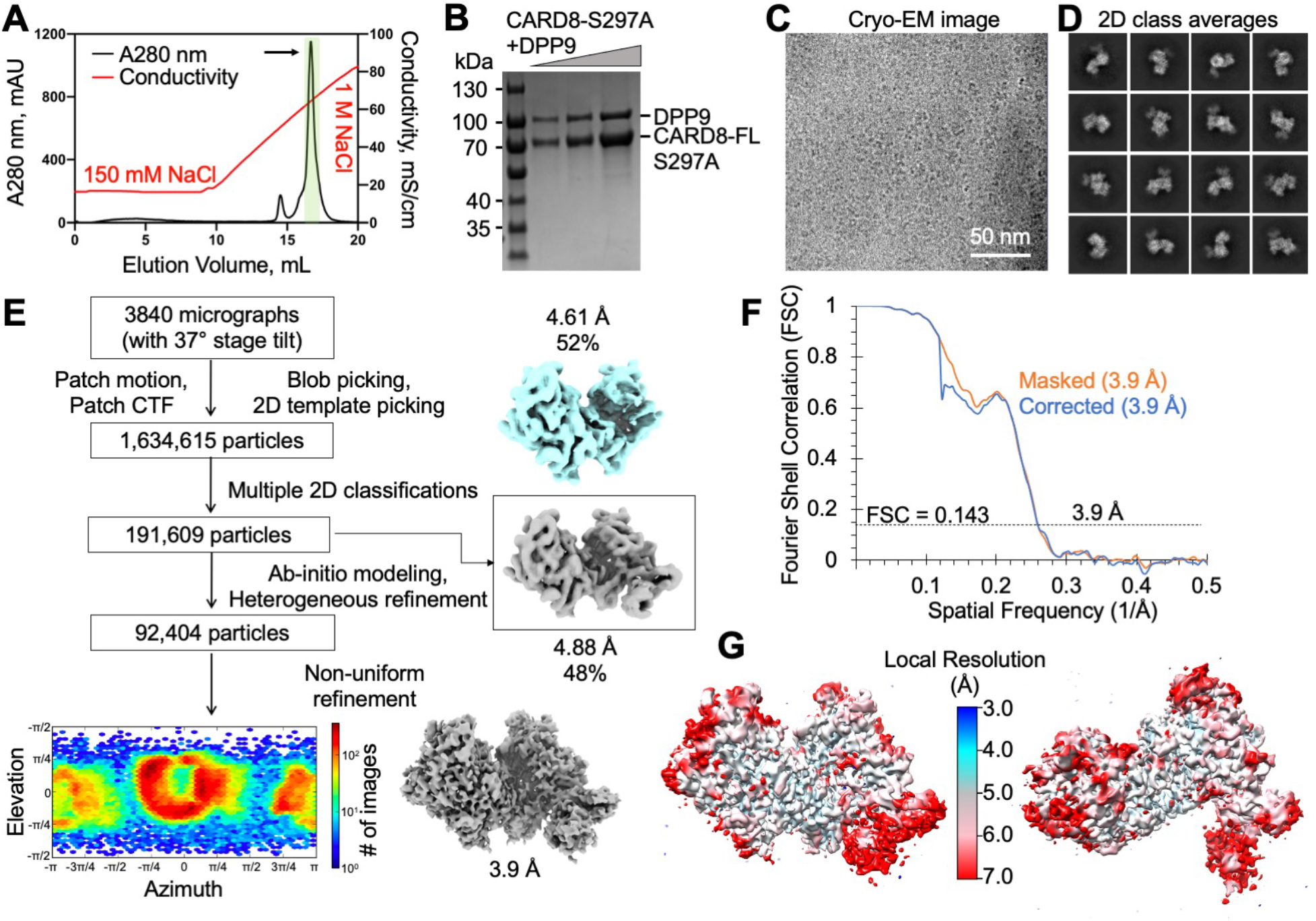
Cryo-EM Structure Determination of the DPP9-CARD8 (S297A) Complex, Related to Figure 2. (A) Purification of the CARD8-DPP9 complex by ion exchange chromatography. (B) SDS-PAGE gel demonstrating autoprocessing deficiency. (C) A representative cryo-EM micrograph. (D) 2D class averages. (E) Workflow for the CARD8-DPP9 complex structure determination with cryoSPARC. (F) Gold-standard Fourier shell correlation (FSC) curves of two independent half maps from cryoSPARC that are either masked (orange) or corrected by noise substitution and masked (blue). (G) Local resolution distribution of the map calculated with cryoSPARC.

**Figure S4.**
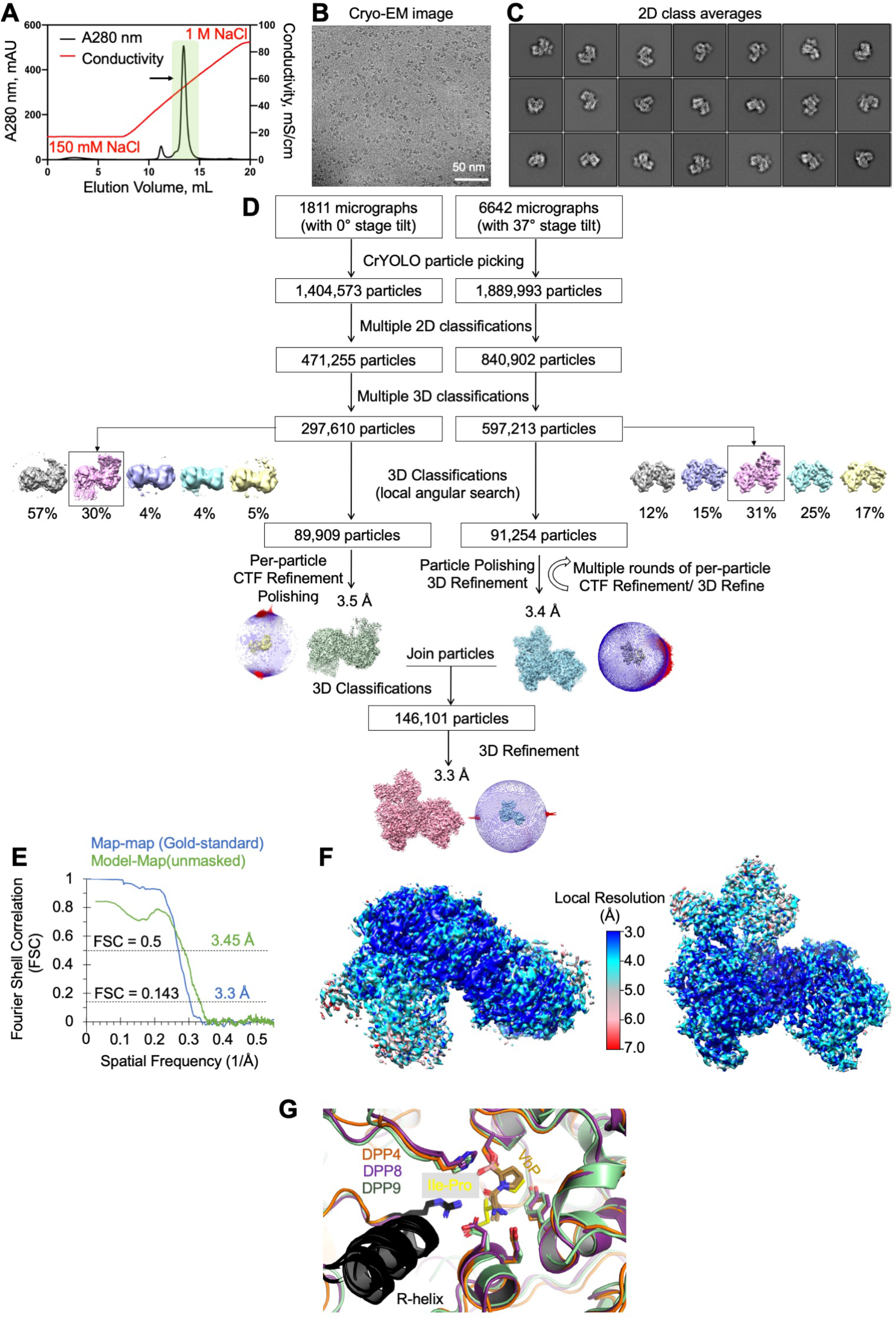
Cryo-EM Structure determination of the DPP9-CARD8-VbP complex, Related to Figure 2. (A) MonoQ anion exchange elution profile of the CARD8-DPP9-VbP complex in running buffer containing 10 μM VbP. An arrow indicates the peak used for subsequent dialysis into a lower salt (150 mM NaCl) buffer and cryo-EM structure determination. (B) A representative cryo-EM micrograph. (C) Representative 2D class averages. (D) Workflow of cryo-EM data analysis conducted with CrYOLO and RELION v3.1. Two datasets, collected at 0° (1811 micrographs) and 37° (6642 micrographs) stage tilts, were combined to yield a structure at 3.3 Å resolution. (E) Gold-standard Fourier Shell Correlation (FSC) curves between the two independent half maps (blue) and between the unmask map and the atomic model (green). (F) Local resolution of the cryo-EM map determined by ResMap and colored as indicated. (G) Structural alignment of the DPP9-CARD8-VbP complex and the published crystal structures of bacterial DPP4 bound to the substrate Ile-Pro (PDB ID: 5YP3) and the DPP8-VbP complex (PDB ID: 6HP8). The VbP molecules (carbon atoms in brown) assume a pose remarkably similar to the model substrate Ile-Pro of DPP4 (carbon atoms in yellow).

**Table S1.**
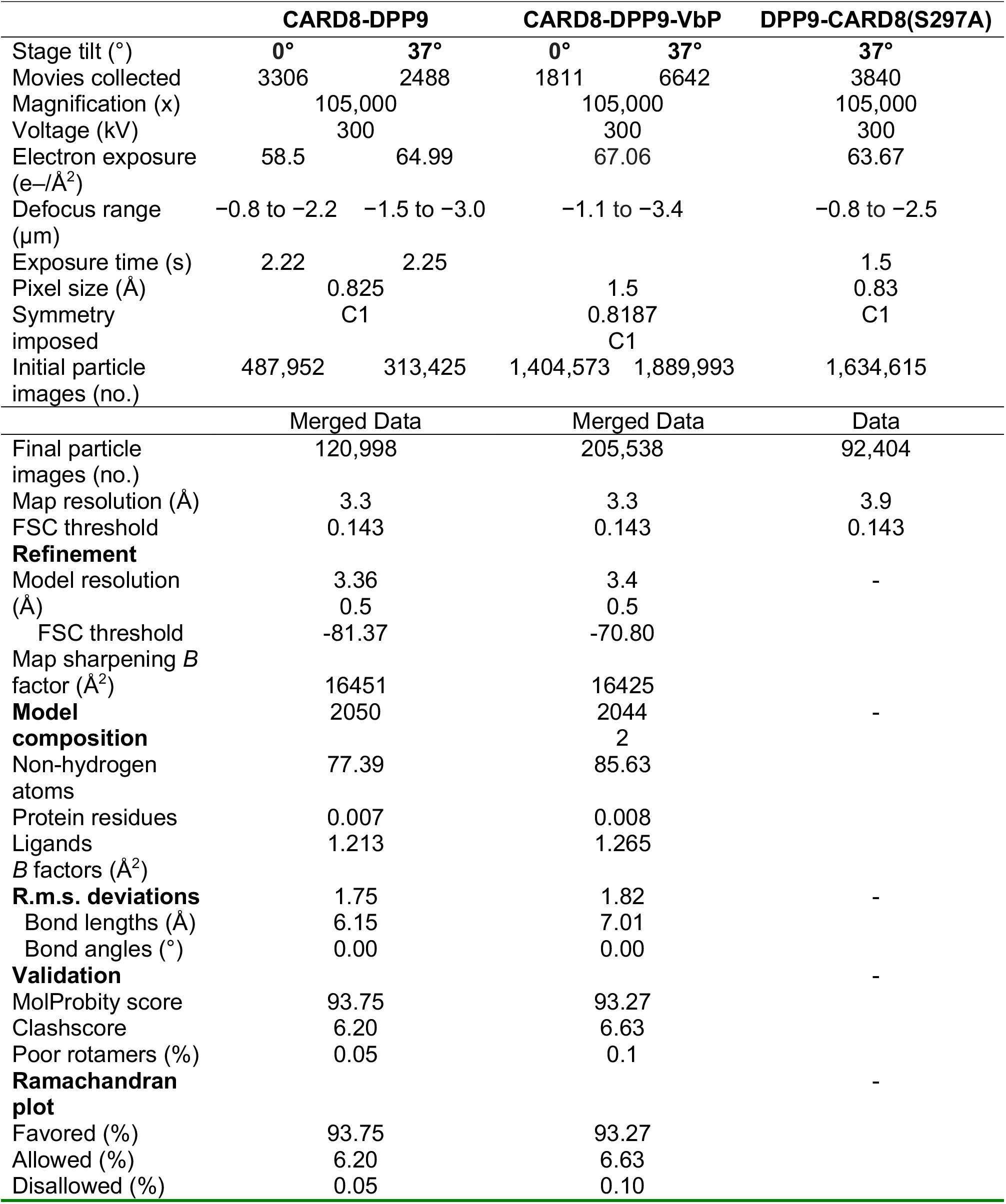
Cryo-EM Data Collection, Refinement, and Validation Statistics, Related to Figures 1 and 2.

